# *KAT2* paralogs prevent dsRNA accumulation and interferon signaling to maintain intestinal stem cells

**DOI:** 10.1101/2023.09.04.556156

**Authors:** Mai-Uyen Nguyen, Sarah Potgieter, Winston Huang, Julie Pfeffer, Sean Woo, Caifeng Zhao, Matthew Lawlor, Richard Yang, Angela Halstead, Sharon Dent, José B. Sáenz, Haiyan Zheng, Zuo-Fei Yuan, Simone Sidoli, Christopher E. Ellison, Michael Verzi

## Abstract

Histone acetyltransferases *KAT2A* and *KAT2B* are paralogs highly expressed in the intestinal epithelium, but their functions are not well understood. In this study, double knockout of murine *Kat2* genes in the intestinal epithelium was lethal, resulting in robust activation of interferon signaling and interferon-associated phenotypes including the loss of intestinal stem cells. Use of pharmacological agents and sterile organoid cultures indicated a cell-intrinsic double-stranded RNA trigger for interferon signaling. Acetyl-proteomics and dsRIP-seq were employed to interrogate the mechanism behind this response, which identified mitochondria-encoded double-stranded RNA as the source of intrinsic interferon signaling. *Kat2a* and *Kat2b* therefore play an essential role in regulating mitochondrial functions as well as maintaining intestinal health.

**Highlights of the work:**

- *Kat2a* and *Kat2b* double knockout in the murine intestinal epithelium triggers activation of the interferon signaling pathway
- *Kat2a/Kat2b* knockout leads to intestinal stem cell loss and other mucosal phenotypes consistent with interferon activation
- Histone PTM mass spec profiling reveals the first in vivo study showing H3K9ac-specific loss with *Kat2a* and *Kat2b* double knockout, yet without correlation to interferon signaling pathway genes
- Comprehensive proteomic analysis identifies non-histone acetyl-lysine targets of KAT2 in the mouse intestine in vivo, including mitochondrial proteins
- Mitochondrial function is compromised upon *Kat2* loss
- dsRIP-seq identifies double-stranded RNA from the mitochondria as a trigger for the intrinsic immune response upon *Kat2* double knockout

## INTRODUCTION

The intestinal epithelium not only serves absorptive functions but also acts as a barrier against pathogens and microbiota within the lumen. Following an infection, nucleotides of pathogenic origin can be detected by pattern recognition sensors (PRRs), such as toll-like receptors (TLRs), stimulator of interferon genes (STING), melanoma differentiation associated gene 5 (MDA5), and retinoic acid-inducible gene I (RIG-I). Besides these pathogen-associated molecular patterns (PAMPs), host-derived danger-associated molecular patterns (DAMPs) from DNA damage and apoptosis can trigger an epithelial response. Host-derived double-stranded RNA (dsRNA) from the nuclear genome via endogenous retrovirus reactivation, or via leaked mitochondrial transcripts, may activate these PRRs^1–9^. In response to detection of these nucleotides, type I and type III interferons (IFNs) are produced in the intestinal epithelium to activate transcription of interferon stimulated genes (ISGs) which mount a defense against threats^10–12^. Furthermore, specialized types of intestinal epithelial cells (IECs) partake in innate immunity, such as Paneth cells via secretion antimicrobial peptides and cytokines^13,14^. Activation of interferon signaling can also have negative consequences on intestinal function, such as with the loss of intestinal stem cells via *Irf2* deletion or poly I:C treatment^15,16^.

*KAT2A* and *KAT2B* encode for proteins General Control Non-derepressible protein 5 (GCN5, also GCN5L2) and p300/CBP-Associated Factor (PCAF), respectively. *KAT2A* and *KAT2B* are paralogous genes and have roughly 70% amino acid sequence homology^17,18^. Both *KAT2A* and *KAT2B* are reported to redundantly catalyze acetylation of lysine residues on histone H3 tails, particularly lysines 9 and 14 (H3K9ac and H3K14ac, respectively). Additionally, these acetyltransferases also catalyze other acetyl and non-acetyl marks^19–32^.

We sought to define the functions of *KAT2* family members in the intestine. In this study, we find that double knockout (DKO) of lysine acetyltransferases 2A (*Kat2a*) and 2B (*Kat2b)* in the intestinal epithelium triggers an intrinsic interferon response and critically compromises intestinal function. Upon inactivation of *Kat2* paralogs in the intestinal epithelium, the IFN pathway is activated, and intestinal stem cells are lost. *Kat2* DKO in IECs leads to compromised mitochondrial function and the accumulation of mitochondrial dsRNA, which serves as the trigger for interferon signaling. As a result, key epithelial functions such as intestinal stem cell renewal are compromised, and the animal can no longer survive.

## METHODS

### Mouse strains

Mice strains are an outbred mix of 129SvEv and C57BL/6J strains. For *Kat2a^flox/flox^*, loxP sites were inserted into the gene locus before exon 3 and after exon 19^33–35^. Whole-body *Kat2b^-/-^* mice with the neomycin construct were generated as previously described^33–35^. *Kat2* mice were provided to our lab by Dr. Sharon Dent (The University of Texas MD Anderson Cancer Center).

### Mouse intestinal epithelium harvesting

Mice taken for harvest were euthanized by cervical dislocation. Mouse intestine was removed and unraveled prior to flushing with PBS, then cut longitudinally. Intestines were further cut into approximately 1-inch pieces, then incubated in 0.5 M ethylenediaminetetraacetic acid (EDTA) in a series of 5, 10, and 25 minute intervals at 4°C on a rotator. After EDTA incubations, intestinal epithelium was isolated by aggressive shaking. For separating crypt and villus preparations, the epithelium-containing EDTA was passed through a 70 micron filter. Isolated crypts and villi were further cleansed in PBS and centrifuged at 4°C for 3 minutes at 200 rpm to make pellets for organoid culture, RNA, and RT-qPCR. Animal protocols are approved by the Rutgers Institutional Animal Care and Use Committee (IACUC). For mouse breeding colonies, mice were fed with special breeder diet (PicoLab Mouse Diet 20, 5058). For weaned mice and other non-breeding adult mice, the standard mouse diet was provided (PicoLab Rodent Diet 20, 5053).

For conditional and inducible knockout of *Kat2a* using intestinal epithelium-specific *Villin-Cre^ERT^*^2^ in mice^36^, 100 mg/kg of tamoxifen (w/v in 10% ethanol and 90% corn oil) was administered by intraperitoneal injections for 4 days daily; for RNA-seq, the same treatment regimen was implemented except for 3 days daily. Control mice received vehicle intraperitoneal injections (10% ethanol and 90% corn oil).

### Intestinal organoid culture

Crypts isolated from mouse intestine were seeded at a density of 250 crypts per 25 µL Matrigel (Cultrex reduced growth factor basement membrane extract, type R1, R&D Systems, Catalog #3433-010-R1) in 48 well plates. Organoids were fed with 250 µL media (N-2, B-27, 0.5 M N-acetylcysteine, 50 µg/mL mouse epidermal growth factor, 100 µg/mL Noggin, R-Spondin [RSPO], and 0.1 mg/mL primocin in advanced DMEM/F-12 supplemented with GlutaMAX, HEPES, and 100 U/mL penicillin/streptomycin) and typically grown until mature and differentiated at 6 to 7 days. For *ex vivo* knockout, 3 to 4 day old primary organoids were treated with either 1 µM tamoxifen in ethanol or an equivalent volume of ethanol vehicle (100% ethanol) for 16 hours, then collected 72 hours later in TRIzol for RNA.

For small molecule inhibitor studies in organoids, all compounds (GSK8612, MedChemExpress Catalog #HY-111941^37^; H-151, MedChemExpress Catalog #HY-112693; ruxolitinib, Cayman Chemical Item #11609) were reconstituted in DMSO. Stock concentrations and dilutions were prepared to attain a 0.2% DMSO concentration across all groups and treatments. DKO was induced on 4-day old primary organoids (tamoxifen treated for 16 hours), with collection in TRIzol for RT-qPCR on day 8 of the same passage. Compound treatment overlapped with DKO induction and was changed every 24 hours for 3 days for a total treatment time of 88 hours. 3 wells of a 48-well plate were collected in 1 mL TRIzol per group and treatment.

For poly dA:dT (InvivoGen, catalog code tlrl-patn) studies in organoids, the compound was reconstituted in water as instructed by the manufacturer. Transfection with poly dA:dT was performed with Lipofectamine 2000 (ThermoFisher, Catalog #11668019) following instructions from the manufacturer for 6 hours on 10 day old primary intestinal organoids, which were separated from Matrigel by pipetting in ice cold PBS. H-151 (2 µM) or 0.2% DMSO vehicle was given concomitantly with poly dA:dT. Concentrations for poly dA:dT was chosen based on manufacturer’s recommendations.

### RNA extraction and cDNA synthesis

For primary mouse intestinal epithelium, approximately 20 to 50 µL of packed epithelial cell pellet was resuspended in 1 mL TRIzol and RNA was isolated according to manufacturer’s instructions. Mouse intestinal epithelium-derived organoids were also resuspended in TRIzol (3 wells from a 48-well plate in 1 mL TRIzol) and extracted for RNA using chloroform. RNA-containing aqueous phase was processed following instructions from Qiagen’s RNeasy micro kit (Qiagen, Catalog #74004). For cDNA synthesis, 500 to 1000 ng of RNA was measured using Nanodrop^TM^ ND-1000 spectrophotometer (ThermoFisher), then processed per the SuperScript III first-strand synthesis protocol provided by Invitrogen (ThermoFisher Scientific, catalog #18080051).

### RT-qPCR

RT-qPCR was conducted with the ABI PRISM 7900HT Sequence Detection System for a 384-well plate loaded with SYBR green (ThermoFisher Scientific, catalog #4309155), cDNA, and reaction mix containing a 0.5 µM final primer concentration. For analysis, C_t_ values were normalized to *Hprt*, *Cyb561d1*, or *Srsf6*, which were chosen due to *Kat2* DKO-induced expression changes in typical housekeeping genes and based on RNA-seq.

### RNA sequencing

whole mouse duodenal epithelial cells were isolated and extracted for RNA using TRIzol (Invitrogen) and chloroform. RNA samples were sent to BGI for transcriptome library preparation, paired-end RNA-sequencing, and quality control check using their DNBSEQ^TM^ platform (40 million reads per sample). Subsequent fastq files were processed using Kallisto (version 0.46.1) with 100 bootstraps for raw read counts and indexed to the mouse (mm9) genome. Transcript abundance HDF5 binary files were imported with tximport, then processed using DESeq2 (version 1.32) for calculating FPKM normalization and for differential expression analysis in R. Genes for analysis were filtered for average FPKM > 1. DESeq2 was used to generate a preranked gene list based on signal-to-noise ratios for conducting gene set enrichment analysis (GSEA)^38^.

### dsRIP-seq

Whole intestinal epithelial cells from the proximal half of the small intestine of 2 control and 2 *Kat2a/Kat2b* DKO mice were collected for dsRIP-seq, following Gao et al^39^. In short, ∼20-30 µL epithelial cell pellet per sample was resuspended in TRIzol (Invitrogen) as inputs. 200 µL pellets per sample were also resuspended in 3 mL of dsRIP buffer [100 mM NaCl, 50 mM Tris-HCl pH 7.4, 3 mM MgCl_2_, 0.5% IGEPAL CA-630 with 1x mammalian ProteaseArrest protease inhibitor (Gbiosciences, Catalog #786-433) and 40 units/mL SUPERase•In™ RNase Inhibitor (ThermoFisher Scientific, Catalog #AM2696) in water] and incubated for 5 minutes on ice, then centrifuged at max speed (13,000 g) for 10 minutes at 4 °C. Supernatant was divided into 3 aliquots for incubation in normal mouse IgG (MilliporeSigma, Catalog #12-371), dsRNA K1 (Cell Signaling Technology #28764), or dsRNA J2 (Cell Signaling Technology #76651) at 5 µL per 300 µL supernatant. Samples in normal mouse IgG, dsRNA K1, or dsRNA J2 were incubated with rotation at 4 °C for 2 hours, then incubated along with pre-washed Protein G Dynabeads (ThermoFisher Scientific, Catalog #10003D), using 25 µL beads per 300 µL supernatant, for 1 hour at 4 °C with rotation. Beads were collected with a magnetic rack and washed four times in a cold room before addition of TRIzol.

RNA extraction for dsRIP was performed using an miRNeasy micro kit (Qiagen, Catalog #217084) following the manufacturer’s protocol and two rounds of chloroform incubations to reduce phenol contamination. RNA was measured using Nanodrop^TM^ ND-1000 spectrophotometer (ThermoFisher) before shipping to Azenta (GeneWiz) for next generation sequencing. Similar to previous reports, IgG samples yielded insufficient amounts of RNA for sequencing and were not included for further analysis^39^. Library prep included rRNA-depleted total RNA followed by cDNA synthesis using random hexamers. FastQC (version 0.12.0) was used for quality checking fastq files. Unique molecular identifiers (UMIs) were provided by Azenta in a separate fastq file. UMIs were linked to their respective sequencing read with umis (version 1.0.3)^40^ and strand-specific reads were aligned to the mm10 genome (from Gencode, GRCm38.p4) using STAR (version 2.7.10b) for splice awareness. The resulting BAM file was then indexed with samtools (version 1.17) and deduplicated with UMI-tools (version 1.1.4)^41–44^. Per-gene raw reads counts were obtained using HTSeq (version 2.0.2) and differential expression analyses were conducted with DESeq2 (version 1.40.1) by calculating ratios of ratios^45,46^. FPKM values were determined using edgeR (version 3.42.2) following the script provided by Gao et al^39,47^. Bigwig files for IGV tracks were generated using BAM files from STAR and deepTools (version 3.5.1)^48^.

### ChIP-seq analysis

Analysis of H3K9ac ChIP-seq from GSE86996^49^ was conducted using Bowtie (v1.3.0)^50^ and deepTools (v3.5.0)^48^ with alignment to the mm9 mouse genome. Positive and negative sense strands were reoriented into the same direction for analysis. Counts from BAM files were used to compare H3K9ac signal from ChIP-seq to the log fold change of FPKM values between *Kat2* DKOs and controls from RNA-seq.

### Histology

Harvested mouse intestine used for paraffin embedding was flushed with PBS and cut longitudinally, then pinned and fixed in 4% paraformaldehyde (PFA) in PBS for 10 minutes prior to Swiss rolling. After overnight incubation in additional 4% PFA, intestine Swiss rolls were briefly rinsed in PBS and dehydrated in a series of 70% to 100% ethanol incubations. Rolls were transferred into xylene for 2 hours, then submerged in melted paraffin for 4 hours and embedded for sectioning. Paraffin embedded tissues were sectioned at a thickness of 5 µm.

### Immunohistochemistry

5 µm paraffin-embedded tissue sections were heated at 65° C and incubated in xylene before processing in increasing concentrations of ethanol. Sections were rehydrated and underwent antigen retrieval with 10 mM sodium citrate buffer, pH 6 along with peroxidase quenching and permeabilized with 0.5% Triton. Samples were blocked in 5% fetal bovine serum (FBS) and incubated overnight with primary antibody diluted in FBS (see Supplementary Table 1 for antibody list). After biotinylated secondary antibody incubation following instructions from VECTASTAIN Elite ABC-HRP-kit (Vector Laboratories, catalog #PK-6101), antigen signal was amplified with avidin/biotin complexing (ABC) for 1 hour and developed with 0.05% 3,’3-diaminobenzidine (DAB, CAS #868272-85-9) and 0.015% hydrogen peroxide in 0.1 M Tris. Tissue was counterstained with hematoxylin, then processed again in increasing ethanol concentrations and xylene prior to mounting.

### Mitochondrial enzymatic activity staining

4 µm sections of unfixed, OCT-embedded jejunal tissue were incubated in NADH-containing buffer (1 mg/mL nitro blue tetrazolium chloride, 5 mM MgCl_2_, 25 mM CoCl_2_, 2 mg/mL NADH, 0.2 M Tris-HCl buffer, pH 7.4) at room temperature until the tissue developed a blue color or buffer containing cytochrome c (5 mg DAB, 10 mg cytochrome c, 20 µg catalase, 750 mg sucrose, 2.5 mL 0.2 M sodium phosphate buffer, pH 7.6 in 7.5 mL deionized water) at 37 °C until the tissue developed a brown color. For mitochondrial complex I activity staining, tissue was then fixed in 4% paraformaldehyde for 10-15 minutes, washed 2 times in water, then mounted in water-based mounting medium^51^. For COX activity staining, tissue was washed in water prior to dehydration with increasing concentrations of ethanol, clearance with xylene, and mounting with a xylene-based medium^52^.

### Cell Lysis and Protein Extraction

Primary intestinal epithelial cells were processed in RIPA buffer (150 mM sodium chloride, 50 mM Tris pH 8.0, 1% NP-40, 0.5% sodium deoxycholate, 0.1% SDS in water) to extract total protein. RIPA buffer was spiked with inhibitors to achieve final concentrations of: 1x protease inhibitor, 1 mM phenylmethylsulfonyl fluoride (PMSF), 1 mM sodium orthovanadate, 10 mM sodium fluoride, and 10 mM sodium butyrate. Samples were sonicated for 2-3 minutes prior to incubating at 4 °C with rotation for 30 minutes. After spin down at max speed in a mini centrifuge for 15 minutes at 4 °C, the resulting supernatant containing protein was saved for additional assays described below. Protein quantification was performed following instructions from the Pierce BCA Protein Assay Kit (ThermoFisher Scientific, catalog #23225), using the SpectraMax M2 (Molecular Devices) plate reader and SoftMax Pro software (Molecular Devices).

### Histone Purification and Post-Translational Modification Mass Spectrometry

Histones were purified from isolated intestinal epithelial cells using acid-based methodology for mass spectrometry of histone post-translational modifications, as described by Sidoli et al and Yuan et al^53,54^. In short, for histone extraction and digestion, histone proteins were extracted from pelleted intestinal epithelial cells to ensure good-quality identification and quantification of single histone marks^53^. Briefly, histones were acid-extracted with chilled 0.2 M sulfuric acid (5:1, sulfuric acid : pellet) and incubated with constant rotation for 4 h at 4°C, followed by precipitation with 33% trichloroacetic acid (TCA) overnight at 4°C. Then, the supernatant was removed, and the tubes were rinsed with ice-cold acetone containing 0.1% HCl, centrifuged and rinsed again using 100% ice-cold acetone. After the final centrifugation, the supernatant was discarded, and the pellet was dried using a vacuum centrifuge. The pellet was dissolved in 50 mM ammonium bicarbonate, pH 8.0, and histones were subjected to derivatization using 5 µL of propionic anhydride and 14 µL of ammonium hydroxide (all Sigma Aldrich) to balance the pH at 8.0. The mixture was incubated for 15 min and the procedure was repeated. Histones were then digested with 1 µg of sequencing grade trypsin (Promega) diluted in 50mM ammonium bicarbonate (1:20, enzyme:sample) overnight at room temperature. Derivatization reaction was repeated to derivatize peptide N-termini. The samples were dried in a vacuum centrifuge.

Prior to mass spectrometry analysis, samples were desalted using a 96-well plate filter (Orochem) packed with 1 mg of Oasis HLB C-18 resin (Waters). Briefly, the samples were resuspended in 100 µl of 0.1% TFA and loaded onto the HLB resin, which was previously equilibrated using 100 µl of the same buffer. After washing with 100 µl of 0.1% TFA, the samples were eluted with a buffer containing 70 µl of 60% acetonitrile and 0.1% TFA and then dried in a vacuum centrifuge.

For LC-MS/MS acquisition and analysis, samples were resuspended in 10 µl of 0.1% TFA and loaded onto a Dionex RSLC Ultimate 300 (Thermo Scientific), coupled online with an Orbitrap Fusion Lumos (Thermo Scientific). Chromatographic separation was performed with a two-column system, consisting of a C-18 trap cartridge (300 µm ID, 5 mm length) and a picofrit analytical column (75 µm ID, 25 cm length) packed in-house with reversed-phase Repro-Sil Pur C18-AQ 3 µm resin. Peptides were separated using a 30 min gradient from 1-30% buffer B (buffer A: 0.1% formic acid, buffer B: 80% acetonitrile + 0.1% formic acid) at a flow rate of 300 nl/min. The mass spectrometer was set to acquire spectra in a data-independent acquisition (DIA) mode. Briefly, the full MS scan was set to 300-1100 m/z in the orbitrap with a resolution of 120,000 (at 200 m/z) and an AGC target of 5x10e5. MS/MS was performed in the orbitrap with sequential isolation windows of 50 m/z with an AGC target of 2x10e5 and an HCD collision energy of 30.

Histone peptides raw files were imported into EpiProfile 2.0 software^54^. From the extracted ion chromatogram, the area under the curve was obtained and used to estimate the abundance of each peptide. In order to achieve the relative abundance of post-translational modifications (PTMs), the sum of all different modified forms of a histone peptide was considered as 100% and the area of the particular peptide was divided by the total area for that histone peptide in all of its modified forms. The relative ratio of two isobaric forms was estimated by averaging the ratio for each fragment ion with different mass between the two species. The resulting peptide lists generated by EpiProfile were exported to Microsoft Excel and further processed for a detailed analysis.

### Acetylomics

Primary intestinal epithelial cells in the proximal half of the small intestine from 4 control and 6 *Kat2a/Kat2b* DKO mice were collected. Protein was extracted with RIPA buffer and processed for acetylomic mass spectrometry as described previously^55^ using PTMScan® HS Acetyl-Lysine Motif (Ac-K) Kit (Cell Signaling Technology #46784).

### Immunoblotting

Protein preps were mixed with loading dye and final concentration of 0.1 M DTT, then heated at 95°C for 10 minutes. For purified histones, 5 µg was loaded for each sample into a NuPAGE^TM^ 4-12% Bis-Tris polyacrylamide gel (ThermoFisher Scientific, catalog #NP0336BOX) and run in NuPAGE^TM^ MES SDS running buffer (ThermoFisher Scientific, catalog #NP0002). Proteins were wet transferred onto PVDF membrane, which was subsequently blocked in 5% bovine serum albumin for phosphorylated proteins or 5% milk for non-phosphorylated proteins for 1 hour at room temperature and incubated overnight with primary antibody in blocking buffer (Supplementary Table 1). After washing in TBS-T, membranes were incubated in secondary HRP antibody for 1 hour. Enhanced chemiluminescence substrate (Ultra Digital-ECL substrate solution, KindleBio #R1002) was used to detect and image protein (KwikQuant Imager, Kindle Biosciences #D1001). Quantification was performed using ImageJ by measuring the mean gray value of each band and subtracting it from the background. Ratios were calculated by dividing the resulting value for the target protein of interest by that of the total protein or control.

### Data accessibility

RNA-seq and dsRIP-seq data from this publication have been deposited to GEO SuperSeries accession number GSE235796 for both RNA-seq (GSE235789) and dsRIP-seq (GSE235794). Mass spectrometry raw files of histone peptides are publicly available on the repository Chorus (https://chorusproject.org/) at the project number 1828. Acetylomics mass spectrometry raw files are available on the repository MassIVE (ftp://MSV000092370@massive.ucsd.edu).

## RESULTS

To study the functions of *KAT2A* and *KAT2B* genes in intestinal health, we generated genetic mouse models which ablate both *Kat2* paralogs in the intestinal epithelium (henceforth referred as *Kat2* DKO). Specifically, we integrated the *Villin-Cre^ERT^*^2^ transgenic allele^36^ with *Kat2a^flox/flox^*alleles^36,56^ and with *Kat2b* germline KO alleles (*Kat2b^-/-^*)^33^. Induction of *Kat2* DKO via tamoxifen treatment of adult mice was efficient, and loss of *Kat2* paralogs was confirmed at the RNA and protein level (Fig. 1a-b, 1f). Induced *Kat2* DKO led to rapid and significant weight loss, with animals typically reaching a humane endpoint and requiring euthanasia within a week after the start of tamoxifen injections (Fig. 1c). These results indicate a vital intestinal function for *Kat2* paralogs.

**Figure 1.**
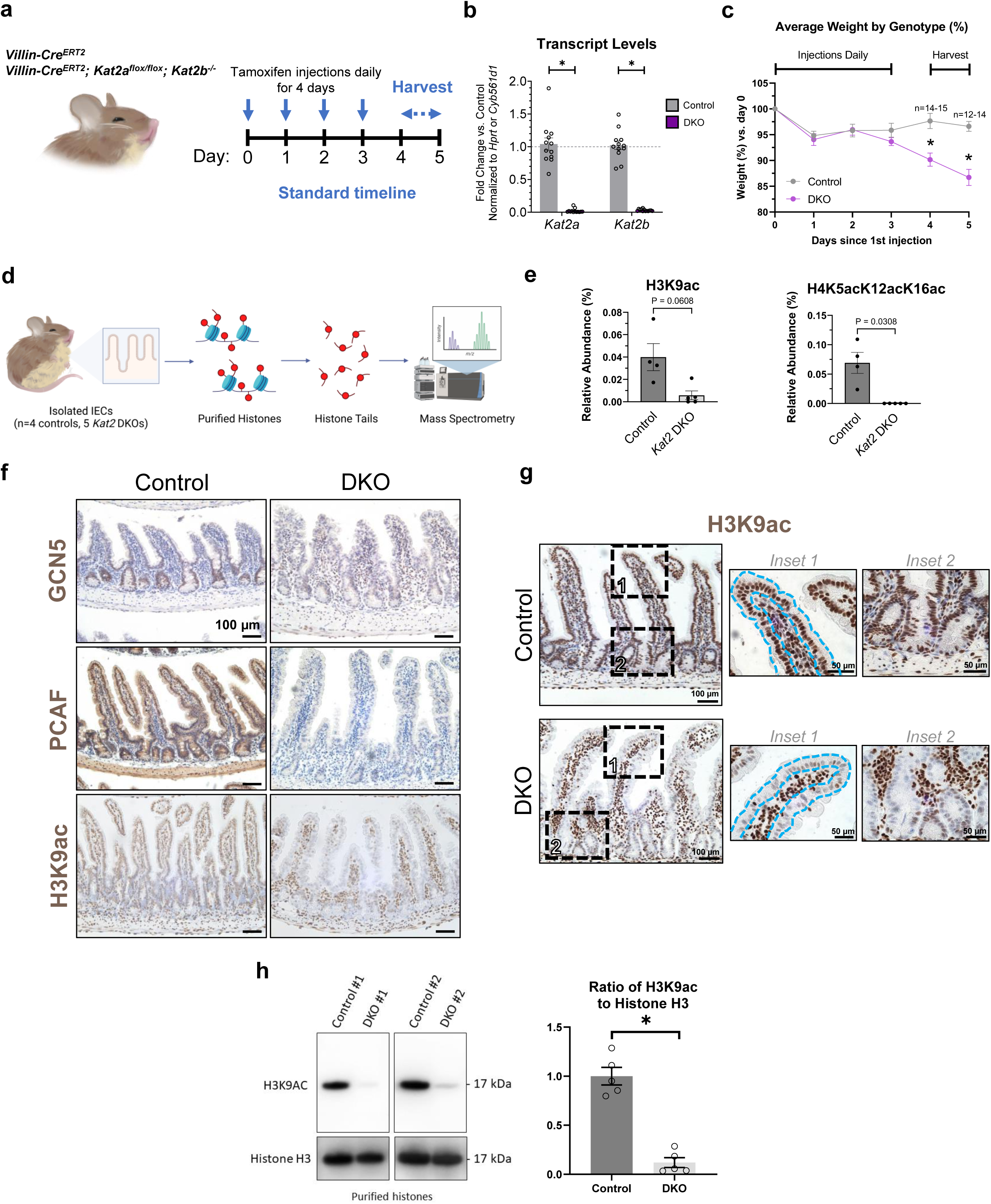
Intestinal epithelium-specific double knockout (DKO) of *Kat2a* and *Kat2b* depletes H3K9ac and is required for survival. (a) Timeline of induction of genetic knockouts and harvest. Vertical arrows indicate the days of tamoxifen injection. Horizontal arrows indicate the average time range for tissue collection. (b) Transcript levels of *Kat2a* and *Kat2b* from *Kat2* DKO mice *in vivo,* normalized to housekeeping gene controls (two-way ANOVA with Bonferroni’s multiple comparisons test, * adj. *p*-val < 0.05, n=12-13/group). (c) Average weights over time in percent, relative to first day of tamoxifen injection, of DKO and control mice show that loss of *Kat2* in the intestine leads to rapid weight loss and death (n=12-15 per group, mixed-effects ANOVA analysis, * adj. *p*-value < 0.05). (d) Schematic of the pipeline for histone PTM mass spectrometry. Created with Biorender.com. (e) Relative abundance (in percent) of H3K9ac and H45acK12acK16ac from histone PTM mass spectrometry of histones from *Kat2* DKO IECs (n=4-5/group, Welch’s *t*-test). (f) Immunohistochemistry of jejunal sections using the indicated antibodies in control and DKO mice (n=3-4 per group), scale bar = 100 µm. Note efficient depletion of GCN5, PCAF, and H3K9ac in DKO tissues. (g) Additional H3K9ac immunostaining; insets outline the intestinal epithelium to highlight the epithelial-specific reduction of H3K9ac immunoreactivity. Scale bar = 100 µm for center images and 50 µm for insets. Immunostaining is representative of 4 biological replicates. (h) Left: immunoblots of two representative samples for H3K9ac and histone H3 expression on purified histones. Right: ratios of H3K9ac to histone H3 in DKOs from densitometric analysis using ImageJ (n=5/group, unpaired *t*-test, * *p*-val < 0.01). All graphs show means with error bars in SEM.

*KAT2A* and *KAT2B* are reported to catalyze histone post-translational modifications (PTMs) H3K9ac and H3K14ac^21,57–61^ but could more broadly affect the histone code. To comprehensively screen changes in histone modifications from *Kat2* DKOs, histone PTM mass spectrometry^53^ was conducted (Fig. 1d). Consistent with previous literature linking *KAT2* to acetylation of histone 3, lysine 9, the relative abundance of H3K9ac in *Kat2* DKO mutants was decreased (Fig. 1e). Interestingly, other changes were observed in histone PTMs, such as nearly complete loss of detection of multiple acetyl-lysine marks on histone H4 involving H4K5ac, H4K8ac, H4K12ac, and H4K16ac in various combinations (Fig. 1e, Supplementary Table 2). Total acetylation across all histone tails did not significantly change, suggesting the specificity of *Kat2a* and *Kat2b* to these specific lysine residues (Supplementary Fig. 1a). Additional changes to histone marks, such as significantly increased methyl marks H3K18me1K23me1 and H3K56me2 (Supplementary Table 2), are not expected to be catalyzed by *Kat2,* which we assume are likely indirect effects of *Kat2* loss. Immunohistochemistry (IHC) confirmed striking depletion of H3K9ac immunoreactivity in the intestinal epithelial nuclei as well as in expanded crypt regions in *Kat2* DKO mice compared to controls (Fig. 1f-g). This loss was not observed in either single *Kat2* KO (Supplementary Fig. 1b-c), suggesting redundant roles of the paralogs in H3K9ac modification. The absence of H3K9ac was additionally quantified by immunoblotting of purified histones from IECs of control and DKO mice (Fig. 1h). Conversely, competing or neighboring marks, such as H3K9me3 and H3K14ac, did not exhibit a qualitative difference between controls and DKO in IHC nor histone PTM mass spectrometry (Supplementary Fig. 1d-e). To our knowledge, this is the first report of H3K9ac absence *in vivo* with a *Kat2* DKO model.

To further investigate the effects of intestinal *Kat2* DKO on intestinal health, immunostaining for stem cell marker OLFM4 and proliferation marker KI67 was conducted (Fig. 2a). Both protein and transcript levels were notably diminished in *Kat2* DKO intestinal epithelium, suggesting *Kat2* function is necessary for maintaining stemness in the intestinal epithelium. To assay for stem cell function, murine intestine epithelial crypts were isolated and cultured as organoids^62,63^. Primary organoids from *in vivo* DKO mice failed to propagate and survive in culture (Fig. 2b), further suggesting compromised stem cell activity upon loss of *Kat2* function.

**Figure 2.**
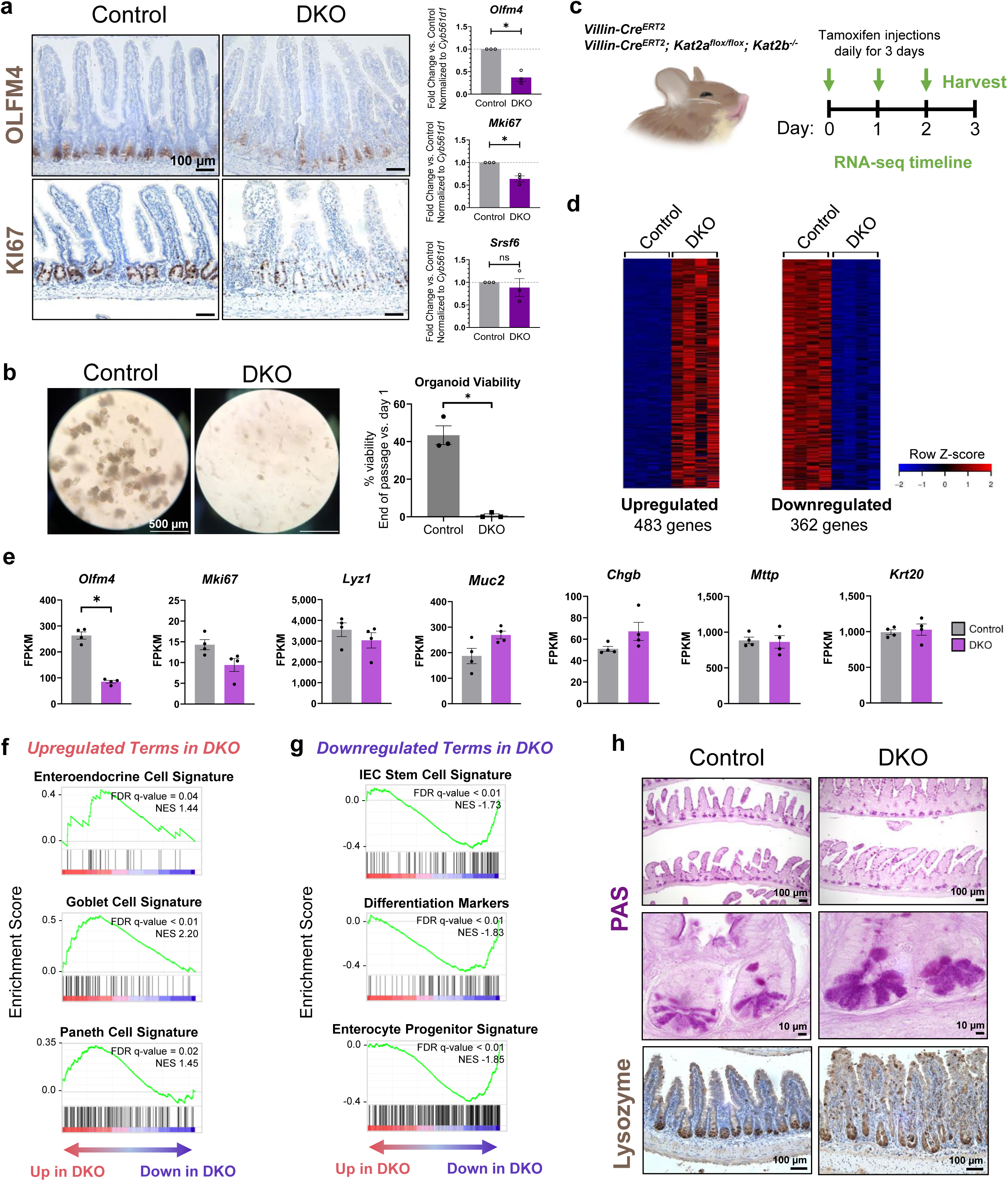
Intestinal epithelium-specific *Kat2a/Kat2b* DKO leads to loss of stem cell renewal and skews expression of distinct intestinal lineage markers. (a) Left: Immunohistochemistry on mouse jejunum for antibodies against OLFM4 and KI67 at 4-7 days after onset of tamoxifen treatment (n=3/group, scale bar = 100 µm). Right: RT-qPCR of *Olfm4* and *Ki67* transcript changes in DKO. *Srsf6* was used as an additional reference gene, and all transcript expression was normalized to housekeeping gene control (n=3/group, unpaired *t*-test, * *p*-val < 0.01). (b) Left: Crypts were collected from mice 4-5 days after the onset of tamoxifen treatment, then seeded in Matrigel for organoid culture. *Ex vivo* primary epithelium-derived intestinal organoids grown for 6 days (n=3/group), scale bar = 500 µm. Right: Survival graph of intestinal organoids (n=3/group, unpaired t-test, * *p*-val < 0.01). (c) Experimental design for RNA-seq. Arrows indicate the days of tamoxifen injection, with tissue collection occurring on day 3. (d) Heatmap of significantly upregulated or downregulated RNA transcripts in *Kat2a/Kat2b* DKO mice (n=4/group, average FPKM > 1, DESeq2 adj. *p*-val < 0.05, fold change >2 or < -2). (e) FPKM values of *Olfm4*, *Mki67*, and select intestinal cell lineage markers from RNA-seq (n=4/group, DESeq2, *\* adj. p*-val < 0.05). GSEA enrichment profiles from RNA-seq that are (f) upregulated or (g) downregulated (n=4/group). (h) PAS staining and IHC for lysozyme of intestinal tissue (n=3/group, scale bar = 10 or 100 µm). All graphs show means with error bars in SEM.

To more comprehensively characterize the functions of *Kat2a* and *Kat2b* in the intestine epithelium, RNA sequencing (RNA-seq) was employed. Epithelium was harvested from control and mutant mice prior to the onset of overt symptoms (Fig. 2c) to understand how the *Kat2* paralogs impact the transcriptome prior to secondary changes of declining mouse health. RNA-seq from *Kat2* DKO IECs and controls resulted in 483 upregulated and 362 downregulated genes (Fig. 2d, average FPKM > 1, fold change > 2 or < -2, DESeq2 adj. *p*-val < 0.05, n=4). Consistent with RT-qPCR data (Fig. 2a), RNA-seq transcript levels for *Olfm4* and *Mki67* decreased with *Kat2* DKO (Fig. 2e). Most other key transcripts used as lineage markers of various intestinal cell types did not significantly change in *Kat2* DKO mutants (Fig. 2e). However, use of broader gene sets associated with intestinal lineages using Gene Set Enrichment Analysis (GSEA)^38^ suggested higher expression of transcripts associated with secretory lineages (enteroendocrine, goblet, and Paneth cell signatures) and reduced expression of transcripts associated with stem cell, differentiation, and enterocyte progenitor signatures^16,64,65^ in *Kat2* DKOs (Fig. 2f-g). The decrease in stem cell transcripts corroborated the lack of organoid forming ability from the DKO crypts (Fig. 2b). Consistent with an increase in Paneth cell gene signature transcripts, *Kat2* DKO mutants also demonstrated increased PAS staining for Paneth cells and disruption in lysozyme protein localization in the intestine (Fig. 2h). Overall, intestinal *KAT2* appears to be required for intestinal stem cell transcript expression and function, and disruption of *KAT2* expression leads to changes in intestinal homeostasis and a rapid decline in the health of the mouse.

To understand how *KAT2* could function to control these aspects of intestinal homeostasis, we further interrogated RNA-seq data. DAVID gene ontology (GO) analysis revealed the most enriched category of genes are associated with viral infection and related immune responses while downregulated genes are associated with oxidoreductase and mitochondrial functions (Fig. 3a-c, Supplementary Table 3). Genes induced after viral infection are often linked to an IFN signaling response. We therefore queried whether the genes induced upon *Kat2* DKO are similar to genes induced upon interferon-lambda (IFN-λ) treatment of intestinal organoids or upon treatment of neonatal mice with interferon-beta (IFN-β) in the intestinal epithelium. We found very robust correlations between these datasets using GSEA (Fig. 3c), indicating that the IFN signaling pathway is activated upon *Kat2* DKO^66^. These findings are consistent with previous literature in which active IFN signaling disturbed intestinal stem cell maintenance and health^15,16^.

**Figure 3.**
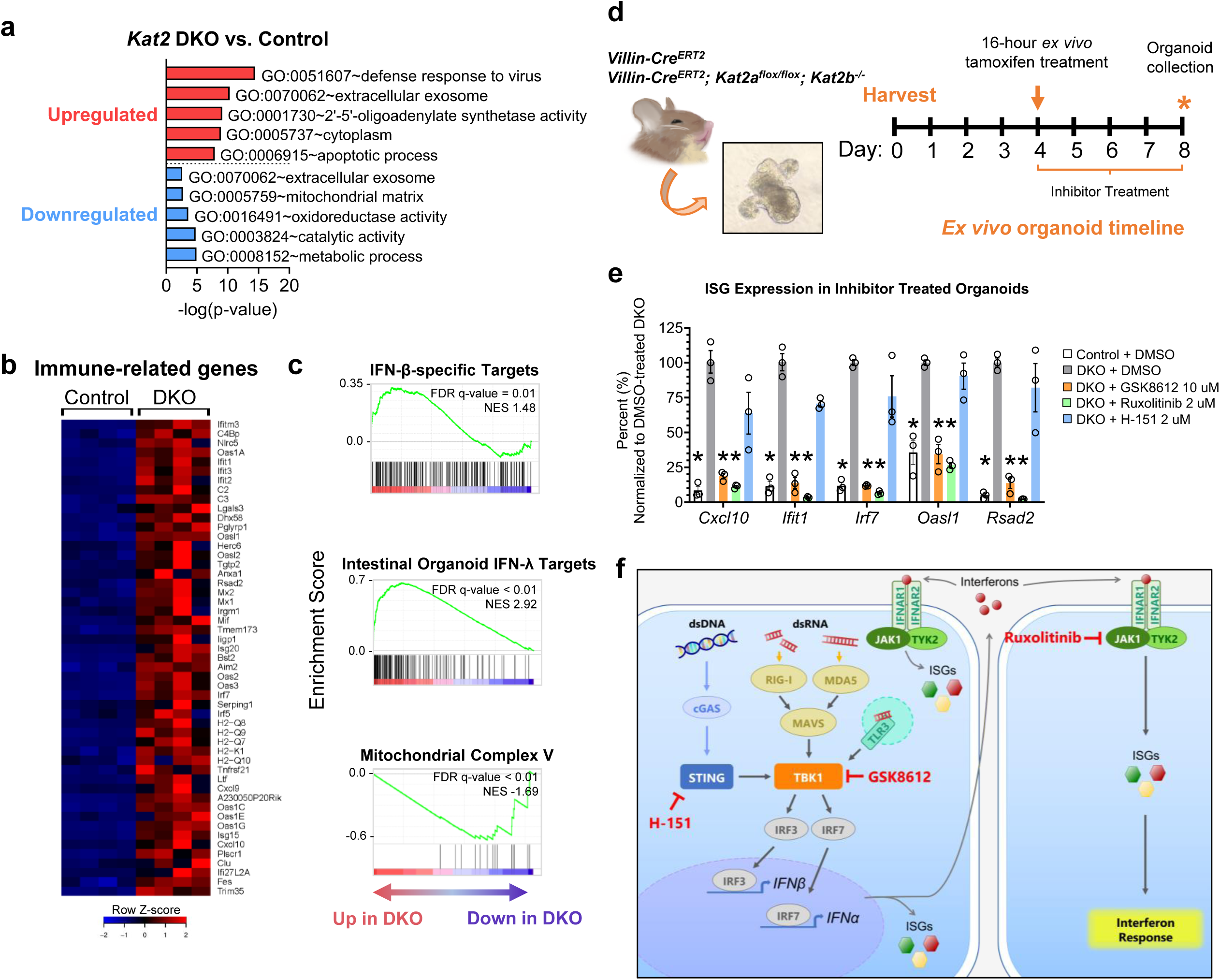
*Kat2a/Kat2b* DKO induces epithelial-intrinsic interferon signaling, which can be suppressed with pharmacological inhibitors of cytoplasmic double-stranded nucleic acid-sensing pathways. (a) Top upregulated (red) and top downregulated (blue) GO terms in DKO samples (483 and 362 genes respectively, from RNA-seq figure 2d) using DAVID GO. (b) Expression profile of immune-related genes populated from RNA-seq and GO analysis (from 5 GO terms: GO:0051607, GO:0009615, GO:0002376, GO:0045087, and GO:0045071). (c) GSEA profiles demonstrate enrichment of IFN-β and IFN-λ targets from GSE142166 and downregulation of mitochondrial complex V in *Kat2a/Kat2b* DKO. (d) Timeline of events for *ex vivo* epithelium-derived intestinal organoid experiments, where mice are harvested on day 0 for 3D organoid culture containing antimicrobials (0.1 mg/mL Primocin, 100 U/mL penicillin-streptomycin). Induction of DKO with 16-hour tamoxifen (1 µM) treatment in *ex vivo* culture begins on day 4. Inhibitor treatment also begins on day 4 and ends on day 8 at the time of organoid collection. (e) RT-qPCR of ISGs from epithelium-derived intestinal organoids with DKO induced in culture, which were treated with pharmacological inhibitors targeting TBK1 (GSK8612, 10 µM), JAK (ruxolitinib, 2 µM), and STING (H-151, 2 µM) for 4 days. Inhibitors were added during and after induction of DKO in culture, with media and inhibitors refreshed daily (n=3 per group, two-way ANOVA with Tukey post-hoc analysis, * adj. *p-*val < 0.05 for the indicated gene versus 0.2% DMSO vehicle-treated DKO, normalized to *Hprt*). Mean shown with error bars in SEM. (f) Schematic of interferon signaling pathway activation via double-stranded nucleic acids. Targets of select inhibitors are shown in red.

Activation of IFN signaling in the *Kat2* DKO mice could be either dependent on a microbial agent or initiated intrinsically in the mouse cells. To discern whether activation of IFN signature genes was dependent upon the microbiome, we induced *Kat2* DKO in the context of established intestinal organoid cultures, which are grown in sterile conditions. Organoids were grown for 4 days in the presence of antimicrobials (0.1 mg/mL primocin) prior to DKO induction (Fig. 3d), and then monitored for ISG expression. Compared to control organoids, DKO organoids exhibited significant elevation of a panel of ISGs (Fig. 3e, Supplementary Fig. 2a). Given the absence of microbial populations and other host-derived cell populations in these epithelial organoid cultures, this interferon response upon *Kat2* DKO appears intrinsic to the intestinal epithelium.

### *Kat2s* suppress a self-derived dsRNA-induced activation of interferon signaling

Cell intrinsic interferon responses can come from several sources, including self-derived double-stranded DNA (dsDNA) and double-stranded RNA (dsRNA). dsDNA and dsRNA sensors initially activate different signaling proteins in the interferon response, but later converge and share the same downstream pathway to trigger transcription of ISGs (Fig. 3f). In the *ex vivo* DKO organoid model, the interferon response can be suppressed using a TBK1-specific inhibitor^37^ (GSK8612) or even more potently suppressed with a JAK inhibitor (ruxolitinib, Fig. 3e). However, inhibition of STING, which is typically activated by dsDNA, failed to repress DKO-induced ISG activation (H-151, Fig. 3e). To confirm efficacy of the STING inhibitor, we transfected wildtype intestinal organoids with a dsDNA mimetic. Poly dA:dT treatment successfully induced ISG expression, as expected, and addition of H-151 effectively repressed this dsDNA response (Supplementary Fig. 2b). Finally, to further test the hypothesis that the *Kat2* response was due to dsRNA detection, we challenged organoids with the dsRNA mimetic, poly I:C. Poly I:C induced the same ISG profile induced by *Kat2* DKOs, and the induction of ISGs by poly I:C could be suppressed with the same pharmacological agents that blocked *Kat2* DKO-induced ISG expression (Supplementary Fig. 2c-f). Thus, dsRNA, and not dsDNA, appears to drive the intrinsic interferon response in *Kat2a/Kat2b* DKO.

### Interferon-stimulated genes are not enriched with nucleosomes containing H3K9ac

To understand whether ISGs are induced in response to dysregulation of transcription associated with loss of H3K9ac in the *Kat2* DKO, we interrogated H3K9ac ChIP-seq from murine jejunal epithelial cells (GSE86996)^49^. H3K9ac signal was aligned at all annotated transcriptional start sites (TSSs) and enriched immediately upstream of the nucleosome-free region associated with the TSS as well as immediately downstream of the TSS, as expected based upon previous profiles of H3K9ac marked histones (GSE86996, Fig. 4a)^49^. K-means clustering resolved 5 distinct H3K9ac patterns at intestinal genes: high signal downstream the TSS (cluster 1), moderate and bidirectional signal from the TSS (cluster 2), moderate signal downstream the TSS (cluster 3), weak signal downstream the TSS (cluster 4), and negligible to no signal (cluster 5, Fig. 4b). We sought to identify what patterns of H3K9ac signal were associated with ISGs differentially expressed in *Kat2* DKO vs. controls from RNA-seq analysis (listed in Fig. 3b). Interestingly, most ISGs that are upregulated in *Kat2* DKO were found in clusters with moderate or low H3K9ac marked promoters (clusters 3-5), suggesting no clear link between ISG induction and epigenomic regulation by *KAT2A-* and *KAT2B-*mediated H3K9ac. ISGs cluster similarly to all DEGs upregulated in *Kat2* DKO (Fig. 4c). DEGs that are downregulated have a slightly greater proportion of genes in clusters 1 and 3 (Fig. 4c). However, DEGs and even all genes from *Kat2* DKO RNA-seq generally do not have a strong correlation with H3K9ac ChIP-seq signal (Fig. 4d), which implies that a non-H3K9ac mechanism is likely activating the interferon response upon *Kat2* DKO.

**Figure 4.**
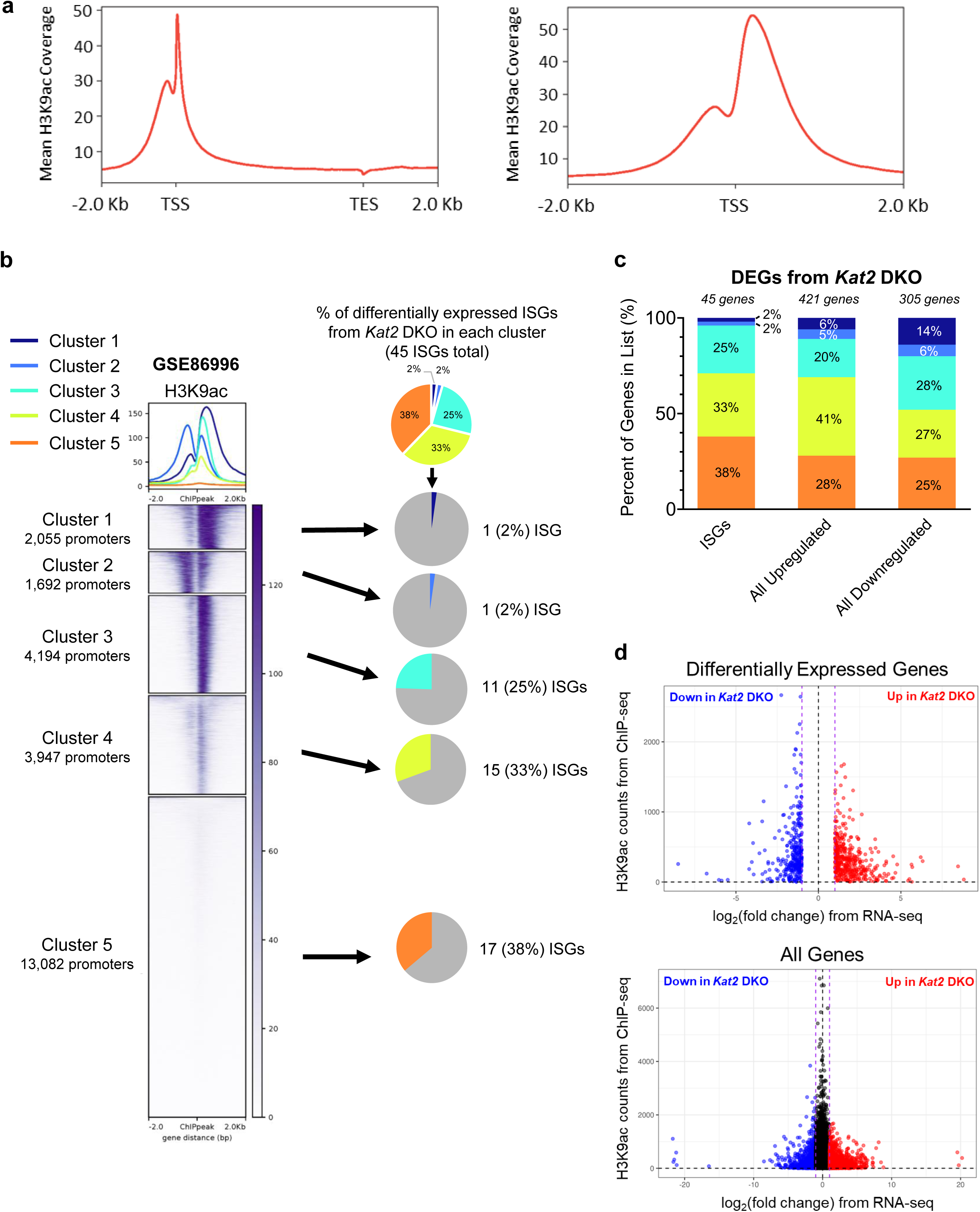
Determination of H3K9ac status at ISGs induced in the *Kat2* DKO. (a) Mean H3K9ac coverage of all murine genes in the jejunal epithelium is shown at a metagene (transcription start site (TSS) to transcription end site (TES) ± 2 Kb (top) and only the TSS ± 2 Kb (bottom)). Positive and negative sense strands were reoriented into the same direction for analysis. Data visuals were generated with deepTools (v3.5.0) using ChIP-seq of H3K9ac from WT mice in GSE86996. (b) Left: K-means clustering of H3K9ac signal within 2 Kb of the TSS forms 5 distinct patterns in the murine jejunal epithelium. Positive and negative sense strands were reoriented into the same direction for analysis. Heatmap visual was generated with deepTools 3.5.0. Right: Pie charts indicating the percentage of upregulated ISGs from *Kat2* DKO RNA-seq belonging to each cluster, highlighting that the presence of H3K9ac in wild type mice does not coincide with interferon stimulated gene expression upon *Kat2* DKO. (c) Bar charts indicating the percentage of ISGs, all DEGs upregulated, and all DEGs downregulated from *Kat2* DKO RNA-seq in each H3K9ac ChIP-seq cluster from Figure 4b. (d) Scatter plots showing the log fold change of *Kat2* DKO FPKM values versus control FPKM values from RNA-seq versus the H3K9ac counts from ChIP-seq (GSE86996) for DEGs (top, from Fig. 4d) or all genes (bottom) from *Kat2* DKO RNA-seq. There does not appear to be a strong correlation between H3K9ac levels and changes in RNA levels in the *Kat2* DKO.

### Identification of *Kat2*-dependent acetylation targets reveals mitochondrial protein acetylation and mitochondrial function are compromised in *Kat2* DKO intestinal epithelium

We therefore aimed to understand whether *KAT2A* and *KAT2B* regulate the interferon response through means beyond transcriptional changes associated with H3K9ac loss. *KAT2A* and *KAT2B* are reported to directly acetylate non-histone proteins in controlling various biological functions in the cell^32,57,67–72^. To investigate what proteins may be under *Kat2*-dependent acetylation, we utilized acetylomic mass spectrometry of intestinal epithelial lysates from control versus *Kat2* DKO mice (Fig. 5a, Supplementary Table 4). In agreement with our findings at the transcript level, the *Kat2* DKO proteome exhibited enrichment in terms related to immune and viral response as well as suppression of oxidoreductase activity (Fig. 5b, Supplementary Table 5). Acetylomic mass spectrometry identified more peptides with diminished acetylation than increased acetylation, as expected upon loss of the *Kat2* acetyltransferases (Fig. 5c). Among the proteins harboring *Kat2*-dependent acetylated residues were those associated with mitochondrial functions (Fig. 5b-c). These included members of the SLC25 family, which are mitochondrial transport carriers of solutes such as ADP/ATP, amino acids, and fatty acids. In light of these changes to the acetylome and diminished mitochondrial transcripts (Fig. 3a) and proteins (Fig. 5b-c), *Kat2* DKO intestinal tissue was stained for activities of enzymes from mitochondrial complex I and IV to assess disruptions in mitochondrial function. Both NADH-oxidation (complex I) and cytochrome c oxidase (COX, complex IV) activities were weaker in *Kat2* DKO (Fig. 5d-e), implying that mitochondrial function is disrupted in the *Kat2* DKO epithelium. These results provide an intriguing look at the acetyl-proteome and link *Kat2* as a potential regulator of mitochondrial function.

**Figure 5.**
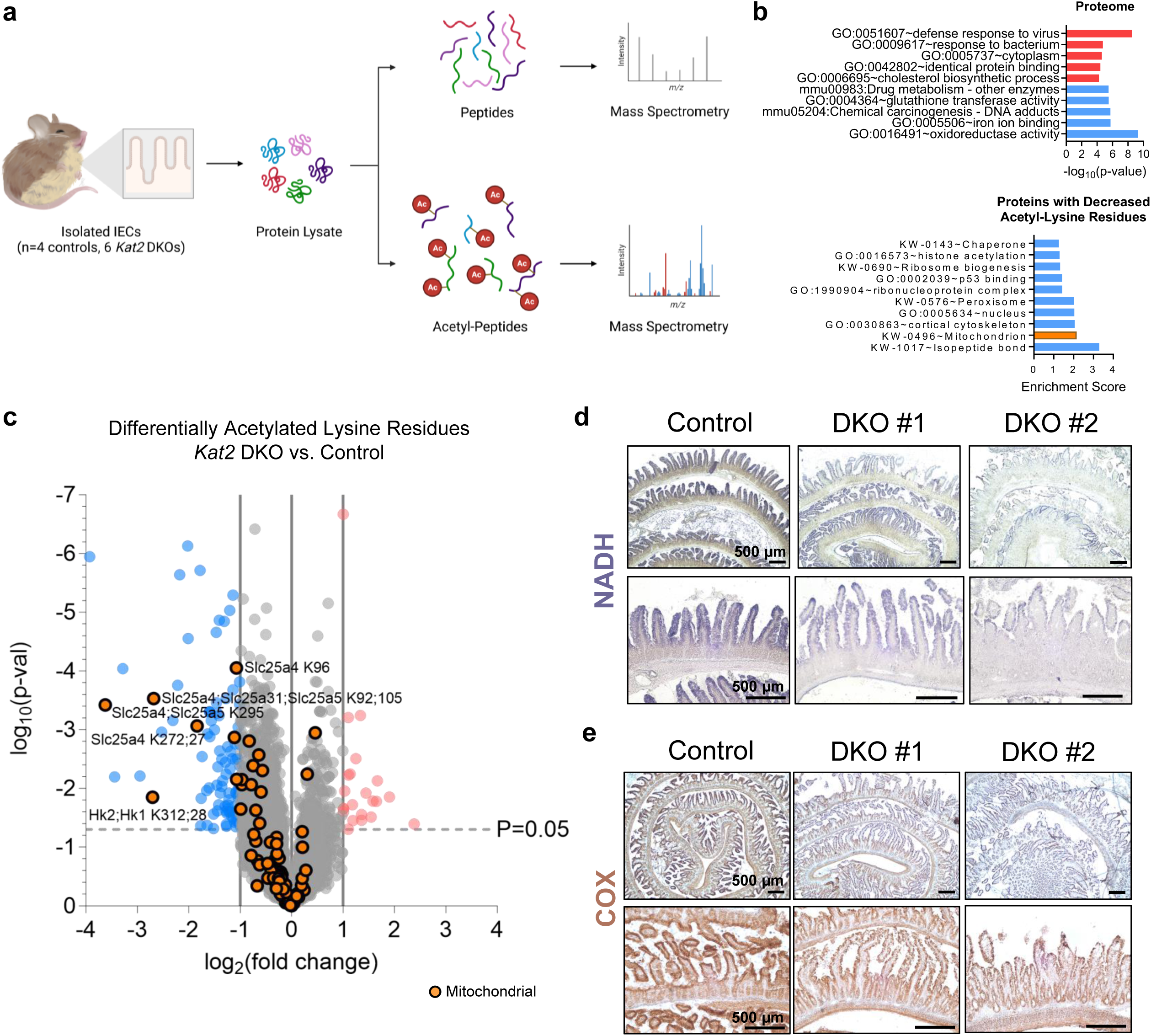
Mitochondrial proteins are differentially acetylated and mitochondrial enzyme activity is compromised in *Kat2* DKO. (a) Schematic of the pipeline for acetylomics mass spectrometry. Created with Biorender.com. (b) GO term analysis of the proteome (top) and non-histone proteins with downregulated acetyl-lysine residues (bottom) from *Kat2* DKO IECs (n=4 controls, 6 DKOs). (c) Volcano plot of acetyl-lysine residues from acetylomics of *Kat2* DKO IECs. Orange points indicate lysine residues on mitochondrial proteins (n=4 controls, 6 DKOs). Stains for activity of mitochondrial enzymes (d) NADH and (e) COX in jejunal intestine sections (n=2-4/group, scale bar = 500 µm).

### Mitochondria-derived dsRNA accumulates in *Kat2* DKO intestinal epithelium

It has previously been demonstrated that compromised mitochondrial function can lead to the generation of mitochondria-derived dsRNA^4,73,74^ and activation of IFN signaling. For example, knockdown of PNPase, an exoribonuclease responsible for degrading mitochondrial transcripts, results in accumulation of cytoplasmic mitochondrial dsRNA and induces transcription of *IFNB* in HeLa cells^4^. Inhibition of ATP synthase from the mitochondrial electron transport chain with oligomycin A also disrupts mitochondrial membrane potential and structure, promoting mitochondrial dsRNA efflux and activating IFN-β and ISGs in chondrosarcoma cells^74^.

To identify the source of dsRNA in *Kat2* DKO epithelium, we conducted sequencing of immunoprecipitated dsRNA (dsRIP-seq) as previously described^39^ using two different monoclonal antibodies designed to recognize dsRNA (anti-dsRNA antibody clones K1 and J2, Fig. 6a, Supplementary Fig. 3). Stranded RNA libraries were generated to allow for detection of both sense and anti-sense transcripts that could contribute to dsRNA generation. Differential enrichment and DAVID GO analyses illustrated an upregulation of dsRNA in dsRIP-seq originating from mitochondria-encoded genes in the *Kat2* DKO samples (Fig. 6b-c, Supplementary Fig. 3a-b, Supplementary Table 6). The highest GO enrichment category of genes associated with dsRNA production are from the mitochondrial genome, regardless of whether the dsRIP-seq was conducted with the K1 or J2 anti-dsRNA antibody. For example, mitochondrial genes *mt-Atp8* and *mt-Nd5* were prominent in dsRNA pulldowns from DKOs but not in control pulldowns nor any input samples (Fig. 6d-e, Supplementary Fig. 3c-d). Notably, *mt-Nd5* and *mt-Nd6* are located on opposite strands at adjacent positions in the mitochondrial genome. These two mitochondrial genes are most susceptible to dsRNA generation due to transcription in opposing directions^75^ and are among the most abundant sources of dsRNA in our dsRIP-seq results (Fig. 6e, Supplementary Fig. 3d). Furthermore, dsRNA pulldowns in DKOs presented with a higher fraction of unique mitochondrial reads (Fig. 6f). Consistent with successful enrichment of dsRNA in pulldown samples, anti-sense transcripts were enriched in the IP samples compared to input RNA (Supplementary Fig. 3e), and total antisense transcripts were higher in mutant samples compared to controls (Fig. 6f). Finally, we note that anti-sense transcript enrichment in the *Kat2* DKOs was specific to the mitochondrial genome, as the nuclear genome had a smaller and consistent fraction of anti-sense reads across all samples (Fig. 6f, Supplementary Fig. 3f). Collectively, these findings imply that *Kat2* DKO-generated dsRNA originates from the mitochondrial genome rather than the nuclear genome, and that mitochondrial dsRNA then triggers an interferon response and subsequent imbalances in intestinal homeostasis.

**Figure 6.**
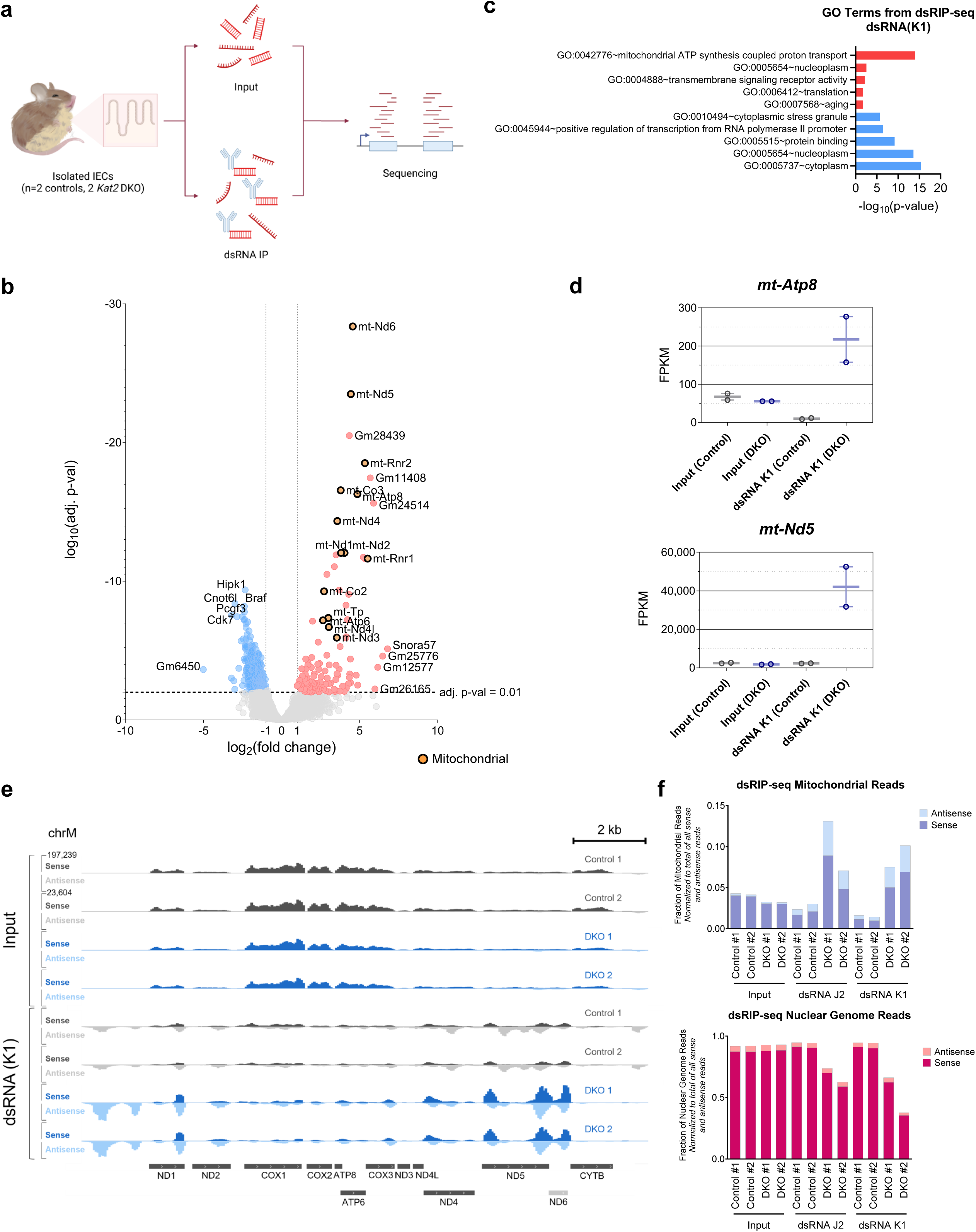
Mitochondrial double stranded RNA (dsRNA) is a likely source of ISG induction in *Kat2a/Kat2b* DKO. (a) Schematic of the workflow for dsRIP-seq. Created with Biorender.com. (b) Volcano plot of dsRIP-seq for dsRNA (K1 antibody) pulldown (n=2/group, DESeq2 adj. *p*-val < 0.01, likelihood ratio test, average FPKM > 1) shows enrichment in dsRNA from mitochondrial transcripts that are specific to anti-dsRNA antibodies in the DKO versus controls and normalized for baseline levels of RNA in input samples. Orange points indicate mitochondrial genes. (c) Top upregulated (red) and top downregulated (blue) GO terms for DKO from dsRIP-seq using dsRNA (K1) antibody and DAVID GO. (d) FPKM values of select dsRNA species upregulated in DKO. Min values, max values, and mean of the values are shown. (e) IGV tracks for input and dsRNA (K1) pulldown samples from dsRIP-seq along the mitochondrial genome. Sense reads are on the same scale (0-197,239) in the upper half of the track and antisense reads on the same scale (0-23,604) in the lower half for each sample. (f) Number of unique mitochondrial (top) or nuclear (bottom) sense or antisense reads from each dsRIP-seq sample, normalized to the total number of all dsRIP-seq reads (including both sense and antisense transcripts).

## DISCUSSION

IEC-specific *Kat2* DKO mutants presented with both an interferon response and stem cell loss. Links between the interferon signaling pathway inhibiting intestinal stemness have been previously documented. Transcription factor IRF2 negatively regulates type I IFN signaling, and deletion of *Irf2* impairs regeneration and maintenance of stem cells in the intestine^15,16^. Furthermore, wildtype colonic organoids from mice treated with poly I:C to induce chronic type I IFN signaling fail to thrive in culture; organoids from *Ifnar1^-/-^* mice undergoing the same treatment, but lacking type I IFN signaling, are capable of growing^15^. It is likely that mice with *Kat2* DKO in IECs follow a similar trajectory from induction of ISGs to suppression in stemness, ultimately leading to the rapid decline in health.

The interferon response is also connected to changes in intestinal homeostasis as the presence of excess interferon signaling induces stem cells to prematurely differentiate into secretory progenitors^16^. An accumulation of immature Paneth cells is observed in *Irf2^-/-^* intestines, with PAS^+^ granule morphology much like in the IEC *Kat2* DKO mutants. Deletion of SETDB1 in the intestine also activates ISGs and results in mislocalization of lysozyme expression^8^.

Abolishment of histone acetyltransferases like *KAT2A* and *KAT2B* and the resulting decrease in histone acetylation might be expected to repress transcription on a global level, but many genes including ISGs were upregulated in IEC-specific *Kat2* DKO mutants. Therefore, *KAT2A* and *KAT2B* may be important components of functioning transcriptional complexes and machinery from initiation to termination, or these histone modulators catalyze acetyl marks on RNA that would confer post-transcriptional stability. For example, *KAT2A* and *KAT2B* have been previously described to acetylate proteins recruited for nucleotide excision repair, such as RPA1, as well as catalyze H3K9ac at sites of DNA damage to mark them for repair^67,76^ In the absence of *KAT2A/KAT2B*, these DNA repair pathways may become inefficient or defective. Future studies should focus their efforts on obtaining a comprehensive understanding of the diverse roles that the *KAT2* genes play both inside and outside the nucleus.

*Kat2* DKO-induced ISG upregulation has been previously documented^57,77^. A study found that knockdown of *KAT2A* and *KAT2B* in human keratinocyte cell lines induced transcription of ISGs at both subconfluence and confluence^77^. In another study, *Kat2* DKO in mouse embryonic fibroblasts also identified upregulation in viral and immune response from RNA-seq, which was attributed to GCN5 directly suppressing TBK1 activity^57^. However, the mechanisms behind this response *in vivo* have not yet been investigated. The intestine-specific knockout in the current study provides new perspectives to explain the mechanism driving this intrinsic interferon response, utilizing novel techniques including acetylomics and dsRIP-seq to elucidate dsRNA from the mitochondria as the instigator in the intestinal epithelium. How *KAT2A* and *KAT2B* modulate mitochondrial function is still uncertain but may relate to acetyl-CoA as a substrate for both post-translational acetylation and oxidative phosphorylation, or the acetylation status of *KAT2*-targeted mitochondrial proteins. Of note, *Kat2* DKO diminished acetylation of solute transporter SLC25 family members at multiple lysines. Modeling simulations of human SLC25A4 determined that acetylation of lysine 23 within the ADP-binding pocket reduced ADP-binding affinity^78^, though this particular residue was not identified in our acetylomics data. While published evidence for other acetyl sites is limited, the SLC25 carrier family is closely tied to mitochondrial pathologies and diseases^79–83^. *Kat2*-catalyzed acetylation may be crucial for conformational changes required for normal carrier functions and providing essential solutes for mitochondrial respiratory complex activity. Notably, formation of mitochondrial dsRNA has been reported to activate type I interferon signaling in HeLa cells upon depletion of polynucleotide phosphorylase PNPase and in patients with hypomorphic mutations in the corresponding *PNPT1* gene^4^. Overall, these mitochondrial defects may cause mitochondrial dsRNA to escape into the cytoplasm and trigger interferon signaling.

## ACKNOWLEDGEMENTS

This research was funded by grants from the New Jersey Commission on Cancer Research (DCHS20PPC023) and NIH (F31CA254086) to M.U.N as well as NIH grants (DK126446 and DK121915) to M.P.V. The Orbitrap Eclipse Tribrid Mass Spectrometer was funded by the NIH (S10OD025140) for H.Z. and C.Z. The work also benefited from the environment at the Cancer Institute of New Jersey (P30CA072720).We thank Jinchuan Xing for key discussions as well as Quan Nguyen and Kiranmayi Vemuri for assisting with bioinformatics.

## SUPPLEMENTARY FIGURES

**Supplementary figure 1.**
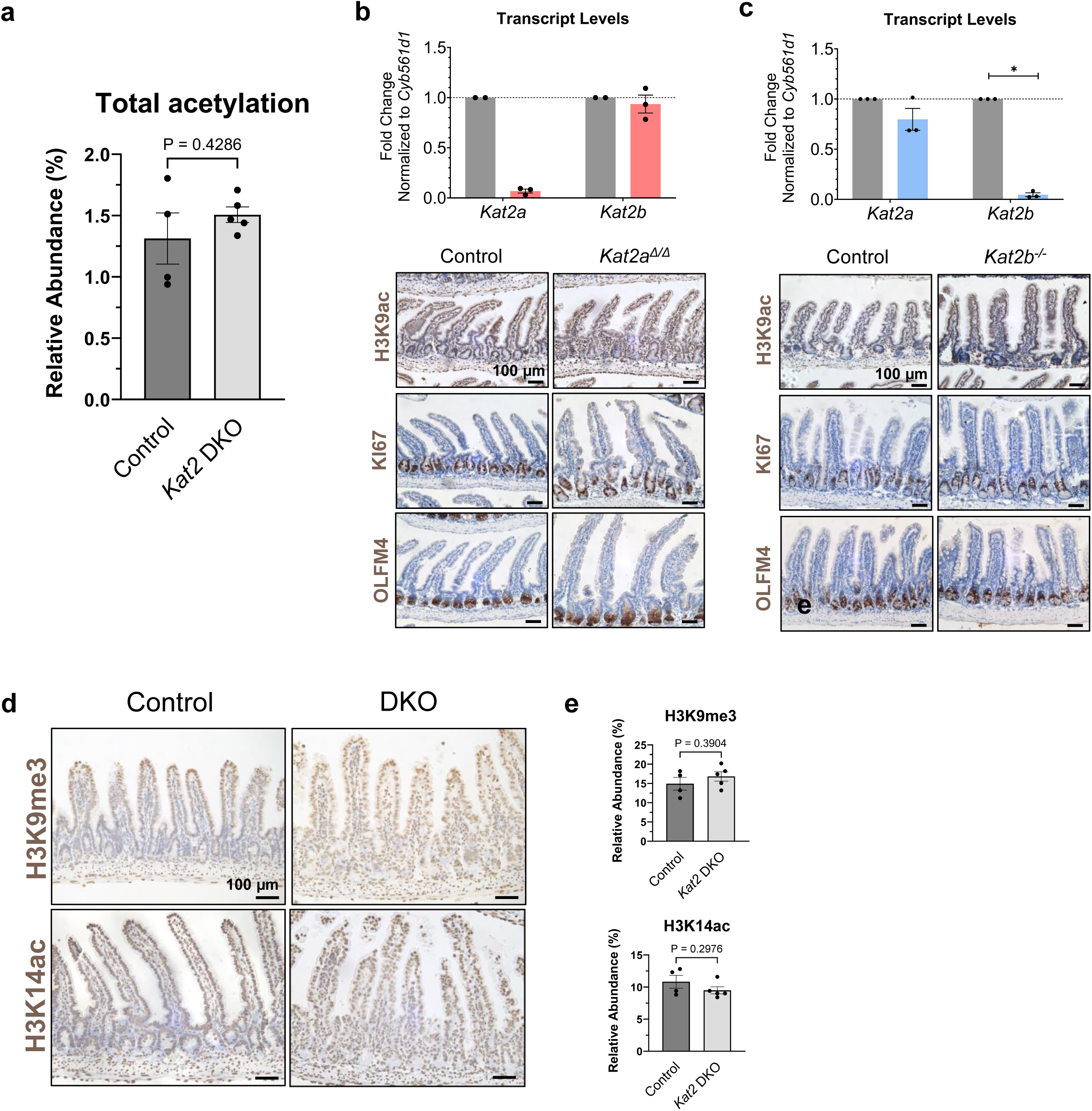
Redundancy between *Kat2* paralogs, and lack of appreciable change in global histone acetylation or specific histone post-translational modifications in the *Kat2* DKO. (a) Relative abundance (in percent) of total acetylation from histone PTM mass spectrometry of histones from *Kat2* DKO IECs (n=4-5/group, Welch’s *t*-test). Mean shown with error bars in SEM. Transcript levels of *Kat2a* and *Kat2b* in IECs (top) and immunostaining of GCN5, H3K9ac, OLFM4, and KI67 (bottom) in (b) IEC-specific *Kat2a^Δ/Δ^* single knockout (n=2-3/group) and (c) *Kat2b^-/-^* single knockout (n=3/group). Scale bar = 100 µm. (d) Immunohistochemistry of jejunal sections against H3K9me3 and H3K14ac in control and DKO mice (n=3-4 per group), scale bar = 100 µm. (e) Relative abundance (in percent) of H3K9me3 and H3K14ac from histone PTM mass spectrometry of histones from *Kat2* DKO IECs (n=4-5/group, Welch’s *t*-test). All graphs show means with error bars in SEM.

**Supplementary figure 2.**
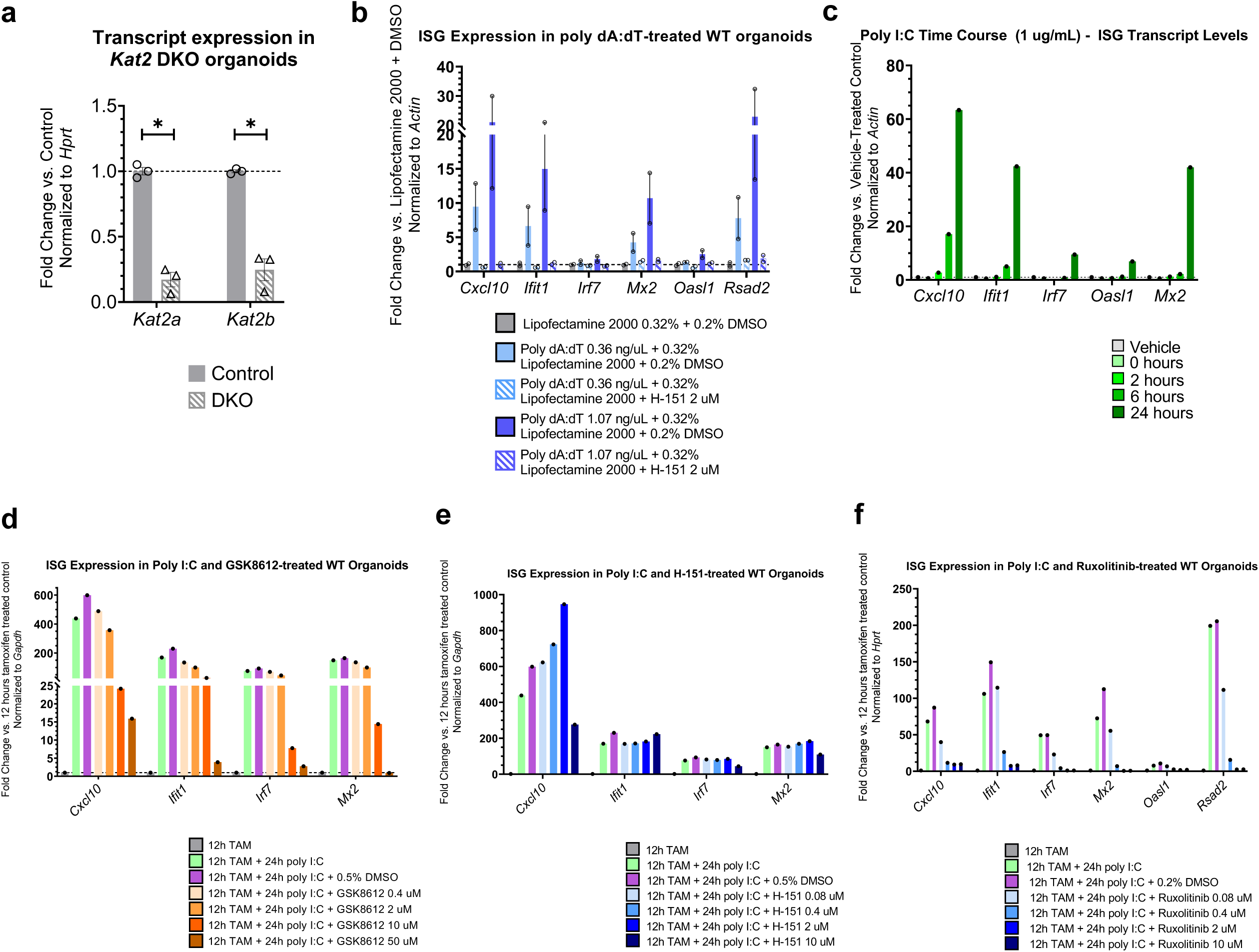
Dissection of how *Kat2* DKO triggers interferon signaling using organoid cultures. (a) Transcript levels of *Kat2a* and *Kat2b* from *Kat2* DKO organoids *ex vivo* normalized to *Hprt* (two-way ANOVA with Bonferroni’s multiple comparisons test, * adj. *p*-val < 0.05, n=3/group). Mean shown with error bars in SEM. (b) Transcript levels of ISGs from 10 day old WT IEC organoid cultures transfected with lipofectamine-complexed poly dA:dT (0.36 or 1.07 ng/µL) and treated with either DMSO vehicle or 2 µM H-151 for 6 hours. Data was normalized to uncomplexed lipofectamine and housekeeping gene *Hprt*. Mean presented with error bars in SEM (n=2/group). These data show efficacy of the H-151 inhibitor, even though H-151 fails to affect the *Kat2* DKO-mediated ISG induction (Fig. 3e). (c) Transcript levels of select ISGs from poly I:C-treated WT IEC organoids at 1 µg/mL for 0, 2, 6, and 24 hours. Data was normalized to water-treated control and *Actin* (n=1/group). (d-f) Organoids were treated with the dsRNA mimetic poly I:C to induce ISGs and test the efficacy of each inhibitor against different components of the interferon response. Transcript levels of select ISGs from WT IEC organoids treated with 1 µg/mL poly I:C in combination with vehicle or (d) GSK8612 at 0.4, 2, 10, or 50 µM, (e) H-151 at 0.08, 0.4, 2, or 10 µM, or (f) ruxolitinib at 0.08, 0.4, 2, or 10 µM for 24 hours. Tamoxifen at 1 µM for 12 hours was added as a control to mimic conditions for IEC organoids with *Kat2* DKO induction in culture. Data was normalized to tamoxifen- and water-treated control as well as *Gapdh* or *Hprt* (n=1/group).

**Supplementary Figure 3.**
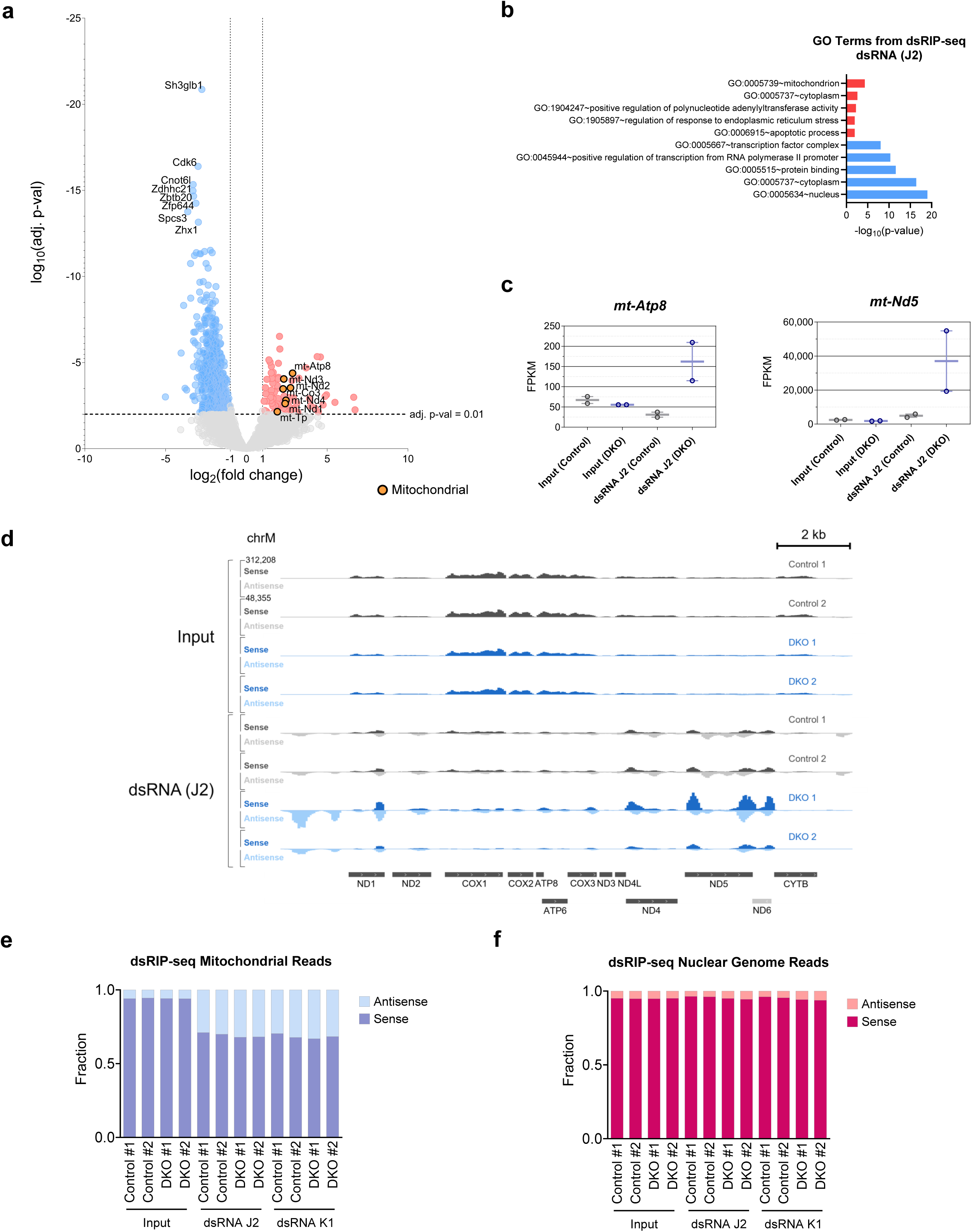
Replicate dsRIP-seq experiment using an independent antibody to enrich for dsRNA also identifies mitochondrial RNA transcripts contributing to dsRNA in *Kat2* DKO. (a) Volcano plot of dsRIP-seq for dsRNA (J2) pulldown (n=2/group, DESeq2 adj. *p*-val < 0.01, likelihood ratio test, average FPKM > 1). Orange points indicate mitochondrial genes. (b) Top upregulated (red) and top downregulated (blue) GO terms for DKO from dsRIP-seq of dsRNA (J2) using DAVID GO. (c) FPKM values of select dsRNA species upregulated in DKO using the dsRNA (J2) antibody. Min values, max values, and mean of the values are shown. (d) IGV tracks for input and dsRNA (J2) pulldown samples from dsRIP-seq along the mitochondrial genome. Sense reads are on the same scale (0-312,208) in the upper half of the track and antisense reads on the same scale (0-48,355) in the lower half for each sample. Fraction of sense and antisense transcripts (right) from each dsRIP-seq sample for the mitochondrial genome (e) and the nuclear genome (f).

## Supplementary Tables

**Supplementary Table 1. List of antibodies and primers used in this publication.**

**Supplementary Table 2. Results of mass spectrometry of histone post-translational modifications from control and *Kat2* DKOs.**

**Supplementary Table 3. DAVID analysis of differentially expressed genes from RNA-seq of *Kat2* DKO.**

**Supplementary Table 4. Results of mass spectrometry of the acetylome from controls and *Kat2* DKOs.**

**Supplementary Table 5. DAVID analysis of proteomic and acetylomic mass spectrometry of *Kat2* DKO.**

**Supplementary Table 6. DAVID analysis of dsRIP-seq (J2 and K1) of Kat2 DKO.**

## REFERENCES

1 Li, T. & Chen, Z. J. The cGAS-cGAMP-STING pathway connects DNA damage to inflammation, senescence, and cancer. J Exp Med 215, 1287–1299 (2018). 10.1084/jem.20180139

2 Yu, L. & Liu, P. Cytosolic DNA sensing by cGAS: regulation, function, and human diseases. Signal Transduct Target Ther 6, 170 (2021). 10.1038/s41392-021-00554-y

3 Sadeq, S., Al-Hashimi, S., Cusack, C. M. & Werner, A. Endogenous Double-Stranded RNA. Noncoding RNA 7 (2021). 10.3390/ncrna7010015

4 Dhir, A. et al. Mitochondrial double-stranded RNA triggers antiviral signalling in humans. Nature 560, 238–242 (2018). 10.1038/s41586-018-0363-0

5 Schlee, M. & Hartmann, G. Discriminating self from non-self in nucleic acid sensing. Nat Rev Immunol 16, 566–580 (2016). 10.1038/nri.2016.78

6 Kariko, K., Buckstein, M., Ni, H. & Weissman, D. Suppression of RNA recognition by Toll-like receptors: the impact of nucleoside modification and the evolutionary origin of RNA. Immunity 23, 165–175 (2005). 10.1016/j.immuni.2005.06.008

7 Wang, R. et al. Gut stem cell necroptosis by genome instability triggers bowel inflammation. Nature 580, 386–390 (2020). 10.1038/s41586-020-2127-x

8 Juznic, L. et al. SETDB1 is required for intestinal epithelial differentiation and the prevention of intestinal inflammation. Gut 70, 485–498 (2021). 10.1136/gutjnl-2020-321339

9 Yuan, H. et al. Lysine catabolism reprograms tumour immunity through histone crotonylation. Nature 617, 818–826 (2023). 10.1038/s41586-023-06061-0

10 Stanifer, M. L., Guo, C., Doldan, P. & Boulant, S. Importance of Type I and III Interferons at Respiratory and Intestinal Barrier Surfaces. Front Immunol 11, 608645 (2020). 10.3389/fimmu.2020.608645

11 Sommereyns, C., Paul, S., Staeheli, P. & Michiels, T. IFN-lambda (IFN-lambda) is expressed in a tissue-dependent fashion and primarily acts on epithelial cells in vivo. PLoS Pathog 4, e1000017 (2008). 10.1371/journal.ppat.1000017

12 Kotenko, S. V. & Durbin, J. E. Contribution of type III interferons to antiviral immunity: location, location, location. J Biol Chem 292, 7295–7303 (2017). 10.1074/jbc.R117.777102

13 Lueschow, S. R. & McElroy, S. J. The Paneth Cell: The Curator and Defender of the Immature Small Intestine. Front Immunol 11, 587 (2020). 10.3389/fimmu.2020.00587

14 Elphick, D. A. & Mahida, Y. R. Paneth cells: their role in innate immunity and inflammatory disease. Gut 54, 1802–1809 (2005). 10.1136/gut.2005.068601

15 Minamide, K. et al. IRF2 maintains the stemness of colonic stem cells by limiting physiological stress from interferon. Sci Rep 10, 14639 (2020). 10.1038/s41598-020-71633-3

16 Sato, T. et al. Regulated IFN signalling preserves the stemness of intestinal stem cells by restricting differentiation into secretory-cell lineages. Nat Cell Biol 22, 919–926 (2020). 10.1038/s41556-020-0545-5

17 Xu, W., Edmondson, D. G. & Roth, S. Y. Mammalian GCN5 and P/CAF acetyltransferases have homologous amino-terminal domains important for recognition of nucleosomal substrates. Mol Cell Biol 18, 5659–5669 (1998). 10.1128/MCB.18.10.5659

18 Nagy, Z. & Tora, L. Distinct GCN5/PCAF-containing complexes function as co-activators and are involved in transcription factor and global histone acetylation. Oncogene 26, 5341–5357 (2007). 10.1038/sj.onc.1210604

19 Huang, H., Sabari, B. R., Garcia, B. A., Allis, C. D. & Zhao, Y. SnapShot: histone modifications. Cell 159, 458–458 e451 (2014). 10.1016/j.cell.2014.09.037

20 Wang, Y. et al. KAT2A coupled with the alpha-KGDH complex acts as a histone H3 succinyltransferase. Nature 552, 273–277 (2017). 10.1038/nature25003

21 Jin, Q. et al. Distinct roles of GCN5/PCAF-mediated H3K9ac and CBP/p300-mediated H3K18/27ac in nuclear receptor transactivation. EMBO J 30, 249–262 (2011). 10.1038/emboj.2010.318

22 Bai, A. H. et al. Dysregulated Lysine Acetyltransferase 2B Promotes Inflammatory Bowel Disease Pathogenesis Through Transcriptional Repression of Interleukin-10. J Crohns Colitis 10, 726–734 (2016). 10.1093/ecco-jcc/jjw020

23 Grant, P. A. et al. Expanded lysine acetylation specificity of Gcn5 in native complexes. J Biol Chem 274, 5895–5900 (1999). 10.1074/jbc.274.9.5895

24 Marmorstein, R. & Zhou, M. M. Writers and readers of histone acetylation: structure, mechanism, and inhibition. Cold Spring Harb Perspect Biol 6, a018762 (2014). 10.1101/cshperspect.a018762

25 Noberini, R. et al. Profiling of Epigenetic Features in Clinical Samples Reveals Novel Widespread Changes in Cancer. Cancers (Basel) 11 (2019). 10.3390/cancers11050723

26 Riss, A. et al. Subunits of ADA-two-A-containing (ATAC) or Spt-Ada-Gcn5-acetyltrasferase (SAGA) Coactivator Complexes Enhance the Acetyltransferase Activity of GCN5. J Biol Chem 290, 28997–29009 (2015). 10.1074/jbc.M115.668533

27 Schiltz, R. L. et al. Overlapping but distinct patterns of histone acetylation by the human coactivators p300 and PCAF within nucleosomal substrates. J Biol Chem 274, 1189–1192 (1999). 10.1074/jbc.274.3.1189

28 Klein, B. J. et al. Recognition of Histone H3K14 Acylation by MORF. Structure 25, 650–654 e652 (2017). 10.1016/j.str.2017.02.003

29 Kuo, M. H. et al. Transcription-linked acetylation by Gcn5p of histones H3 and H4 at specific lysines. Nature 383, 269–272 (1996). 10.1038/383269a0

30 Leemhuis, H., Packman, L. C., Nightingale, K. P. & Hollfelder, F. The human histone acetyltransferase P/CAF is a promiscuous histone propionyltransferase. Chembiochem 9, 499–503 (2008). 10.1002/cbic.200700556

31 Cieniewicz, A. M. et al. The bromodomain of Gcn5 regulates site specificity of lysine acetylation on histone H3. Mol Cell Proteomics 13, 2896–2910 (2014). 10.1074/mcp.M114.038174

32 Xia, L. et al. Modulation of IL-6 Expression by KLF4-Mediated Transactivation and PCAF-Mediated Acetylation in Sublytic C5b-9-Induced Rat Glomerular Mesangial Cells. Front Immunol 12, 779667 (2021). 10.3389/fimmu.2021.779667

33 Xu, W. et al. Loss of Gcn5l2 leads to increased apoptosis and mesodermal defects during mouse development. Nat Genet 26, 229–232 (2000). 10.1038/79973

34 Yamauchi, T. et al. Distinct but overlapping roles of histone acetylase PCAF and of the closely related PCAF-B/GCN5 in mouse embryogenesis. Proc Natl Acad Sci U S A 97, 11303–11306 (2000). 10.1073/pnas.97.21.11303

35 Lin, W. et al. Developmental potential of Gcn5(-/-) embryonic stem cells in vivo and in vitro. Dev Dyn 236, 1547–1557 (2007). 10.1002/dvdy.21160

36 el Marjou, F., et al. Tissue-specific and inducible Cre-mediated recombination in the gut epithelium. Genesis 39, 186–193 (2004). 10.1002/gene.20042

37 Thomson, D. W. et al. Discovery of GSK8612, a Highly Selective and Potent TBK1 Inhibitor. ACS Med Chem Lett 10, 780–785 (2019). 10.1021/acsmedchemlett.9b00027

38 Subramanian, A. et al. Gene set enrichment analysis: a knowledge-based approach for interpreting genome-wide expression profiles. Proc Natl Acad Sci U S A 102, 15545–15550 (2005). 10.1073/pnas.0506580102

39 Gao, Y., Chen, S., Halene, S. & Tebaldi, T. Transcriptome-wide quantification of double-stranded RNAs in live mouse tissues by dsRIP-Seq. STAR Protoc 2, 100366 (2021). 10.1016/j.xpro.2021.100366

40 Svensson, V. et al. Power analysis of single-cell RNA-sequencing experiments. Nat Methods 14, 381–387 (2017). 10.1038/nmeth.4220

41 Frankish, A. et al. Gencode 2021. Nucleic Acids Res 49, D916–D923 (2021). 10.1093/nar/gkaa1087

42 Dobin, A. et al. STAR: ultrafast universal RNA-seq aligner. Bioinformatics 29, 15–21 (2013). 10.1093/bioinformatics/bts635

43 Li, H. et al. The Sequence Alignment/Map format and SAMtools. Bioinformatics 25, 2078–2079 (2009). 10.1093/bioinformatics/btp352

44 Smith, T., Heger, A. & Sudbery, I. UMI-tools: modeling sequencing errors in Unique Molecular Identifiers to improve quantification accuracy. Genome Res 27, 491–499 (2017). 10.1101/gr.209601.116

45 Anders, S., Pyl, P. T. & Huber, W. HTSeq--a Python framework to work with high-throughput sequencing data. Bioinformatics 31, 166–169 (2015). 10.1093/bioinformatics/btu638

46 Love, M. I., Huber, W. & Anders, S. Moderated estimation of fold change and dispersion for RNA-seq data with DESeq2. Genome Biol 15, 550 (2014). 10.1186/s13059-014-0550-8

47 Robinson, M. D., McCarthy, D. J. & Smyth, G. K. edgeR: a Bioconductor package for differential expression analysis of digital gene expression data. Bioinformatics 26, 139–140 (2010). 10.1093/bioinformatics/btp616

48 Ramirez, F. et al. deepTools2: a next generation web server for deep-sequencing data analysis. Nucleic Acids Res 44, W160–165 (2016). 10.1093/nar/gkw257

49 Chen, S. et al. Intestinal NCoR1, a regulator of epithelial cell maturation, controls neonatal hyperbilirubinemia. Proc Natl Acad Sci U S A 114, E1432–E1440 (2017). 10.1073/pnas.1700232114

50 Langmead, B., Trapnell, C., Pop, M. & Salzberg, S. L. Ultrafast and memory-efficient alignment of short DNA sequences to the human genome. Genome Biol 10, R25 (2009). 10.1186/gb-2009-10-3-r25

51 Christensen, T. & Diemer, N. H. Reduction of mitochondrial electron transport complex activity is restricted to the ischemic focus after transient focal cerebral ischemia in rats: a histochemical volumetric analysis. Neurochem Res 28, 1805–1812 (2003). 10.1023/a:1026111506307

52 Hench, J. et al. A tissue-specific approach to the analysis of metabolic changes in Caenorhabditis elegans. PLoS One 6, e28417 (2011). 10.1371/journal.pone.0028417

53 Sidoli, S., Bhanu, N. V., Karch, K. R., Wang, X. & Garcia, B. A. Complete Workflow for Analysis of Histone Post-translational Modifications Using Bottom-up Mass Spectrometry: From Histone Extraction to Data Analysis. J Vis Exp (2016). 10.3791/54112

54 Yuan, Z. F. et al. EpiProfile 2.0: A Computational Platform for Processing Epi-Proteomics Mass Spectrometry Data. J Proteome Res 17, 2533–2541 (2018). 10.1021/acs.jproteome.8b00133

55 Lancho, O. et al. A Therapeutically Targetable NOTCH1-SIRT1-KAT7 Axis in T-cell Leukemia. Blood Cancer Discov 4, 12–33 (2023). 10.1158/2643-3230.BCD-22-0098

56 Lin, W. et al. Proper expression of the Gcn5 histone acetyltransferase is required for neural tube closure in mouse embryos. Dev Dyn 237, 928–940 (2008). 10.1002/dvdy.21479

57 Jin, Q. et al. Gcn5 and PCAF negatively regulate interferon-beta production through HAT-independent inhibition of TBK1. EMBO Rep 15, 1192–1201 (2014). 10.15252/embr.201438990

58 Sen, R. et al. Kat2a and Kat2b Acetyltransferase Activity Regulates Craniofacial Cartilage and Bone Differentiation in Zebrafish and Mice. J Dev Biol 6 (2018). 10.3390/jdb6040027

59 Tjeertes, J. V., Miller, K. M. & Jackson, S. P. Screen for DNA-damage-responsive histone modifications identifies H3K9Ac and H3K56Ac in human cells. EMBO J 28, 1878–1889 (2009). 10.1038/emboj.2009.119

60 Ravnskjaer, K. et al. Glucagon regulates gluconeogenesis through KAT2B- and WDR5-mediated epigenetic effects. J Clin Invest 123, 4318–4328 (2013). 10.1172/JCI69035

61 Han, Y. et al. Histone acetylation and histone acetyltransferases show significant alterations in human abdominal aortic aneurysm. Clin Epigenetics 8, 3 (2016). 10.1186/s13148-016-0169-6

62 Sato, T. & Clevers, H. Primary mouse small intestinal epithelial cell cultures. Methods Mol Biol 945, 319–328 (2013). 10.1007/978-1-62703-125-7_19

63 Sato, T. et al. Single Lgr5 stem cells build crypt-villus structures in vitro without a mesenchymal niche. Nature 459, 262–265 (2009). 10.1038/nature07935

64 Merlos-Suarez, A. et al. The intestinal stem cell signature identifies colorectal cancer stem cells and predicts disease relapse. Cell Stem Cell 8, 511–524 (2011). 10.1016/j.stem.2011.02.020

65 Haber, A. L. et al. A single-cell survey of the small intestinal epithelium. Nature 551, 333–339 (2017). 10.1038/nature24489

66 Van Winkle, J. A., Constant, D. A., Li, L. & Nice, T. J. Selective Interferon Responses of Intestinal Epithelial Cells Minimize Tumor Necrosis Factor Alpha Cytotoxicity. J Virol 94 (2020). 10.1128/JVI.00603-20

67 Zhao, M. et al. PCAF/GCN5-Mediated Acetylation of RPA1 Promotes Nucleotide Excision Repair. Cell Rep 20, 1997–2009 (2017). 10.1016/j.celrep.2017.08.015

68 Ghosh, T. K. et al. Acetylation of TBX5 by KAT2B and KAT2A regulates heart and limb development. J Mol Cell Cardiol 114, 185–198 (2018). 10.1016/j.yjmcc.2017.11.013

69 Fournier, M. & Tora, L. KAT2-mediated PLK4 acetylation contributes to genomic stability by preserving centrosome number. Mol Cell Oncol 4, e1270391 (2017). 10.1080/23723556.2016.1270391

70 Fournier, M. et al. KAT2A/KAT2B-targeted acetylome reveals a role for PLK4 acetylation in preventing centrosome amplification. Nat Commun 7, 13227 (2016). 10.1038/ncomms13227

71 Bondy-Chorney, E., Denoncourt, A., Sai, Y. & Downey, M. Nonhistone targets of KAT2A and KAT2B implicated in cancer biology (1). Biochem Cell Biol 97, 30–45 (2019). 10.1139/bcb-2017-0297

72 Cai, K. et al. C5a promotes the proliferation of human nasopharyngeal carcinoma cells through PCAF-mediated STAT3 acetylation. Oncol Rep 32, 2260–2266 (2014). 10.3892/or.2014.3420

73 Chen, Y. G. & Hur, S. Cellular origins of dsRNA, their recognition and consequences. Nat Rev Mol Cell Biol (2021). 10.1038/s41580-021-00430-1

74 Kim, S. et al. Mitochondrial double-stranded RNAs govern the stress response in chondrocytes to promote osteoarthritis development. Cell Rep 40, 111178 (2022). 10.1016/j.celrep.2022.111178

75 Mercer, T. R. et al. The human mitochondrial transcriptome. Cell 146, 645–658 (2011). 10.1016/j.cell.2011.06.051

76 Guo, R., Chen, J., Mitchell, D. L. & Johnson, D. G. GCN5 and E2F1 stimulate nucleotide excision repair by promoting H3K9 acetylation at sites of damage. Nucleic Acids Res 39, 1390–1397 (2011). 10.1093/nar/gkq983

77 Walters, B. W. et al. Divergent functions of histone acetyltransferases KAT2A and KAT2B in keratinocyte self-renewal and differentiation. J Cell Sci (2023). 10.1242/jcs.260723

78 Mielke, C. et al. Adenine nucleotide translocase is acetylated in vivo in human muscle: Modeling predicts a decreased ADP affinity and altered control of oxidative phosphorylation. Biochemistry 53, 3817–3829 (2014). 10.1021/bi401651e

79 Kunji, E. R. S., King, M. S., Ruprecht, J. J. & Thangaratnarajah, C. The SLC25 Carrier Family: Important Transport Proteins in Mitochondrial Physiology and Pathology. Physiology (Bethesda) 35, 302–327 (2020). 10.1152/physiol.00009.2020

80 Palmieri, F. & Monne, M. Discoveries, metabolic roles and diseases of mitochondrial carriers: A review. Biochim Biophys Acta 1863, 2362–2378 (2016). 10.1016/j.bbamcr.2016.03.007

81 Chen, Y. J. et al. An integrated bioinformatic investigation of mitochondrial solute carrier family 25 (SLC25) in colon cancer followed by preliminary validation of member 5 (SLC25A5) in tumorigenesis. Cell Death Dis 13, 237 (2022). 10.1038/s41419-022-04692-1

82 Iacobazzi, V. et al. Mitochondrial carriers in inflammation induced by bacterial endotoxin and cytokines. Biol Chem 398, 303–317 (2017). 10.1515/hsz-2016-0260

83 Mosaoa, R., Kasprzyk-Pawelec, A., Fernandez, H. R. & Avantaggiati, M. L. The Mitochondrial Citrate Carrier SLC25A1/CIC and the Fundamental Role of Citrate in Cancer, Inflammation and Beyond. Biomolecules 11 (2021). 10.3390/biom11020141

